# Multi-sample SPIM image acquisition, processing and analysis of vascular growth in zebrafish

**DOI:** 10.1101/478149

**Authors:** Stephan Daetwyler, Ulrik Günther, Carl D. Modes, Kyle Harrington, Jan Huisken

## Abstract

To quantitatively understand biological processes that occur over many hours or days, it is desirable to image multiple samples simultaneously and automatically process and analyze the resulting datasets. Here, we present a complete multi-sample preparation, imaging, processing, and analysis workflow to determine the development of the vascular volume in zebrafish. Up to five live embryos were mounted and imaged simultaneously over several days using selective plane illumination microscopy (SPIM). The resulting large imagery dataset of several terabytes was processed in an automated manner on a high-performance computer cluster and segmented with a novel segmentation approach that uses images of red blood cells as training data. This analysis yielded a precise quantification of growth characteristics of the whole vascular network, head vasculature, and tail vasculature over development. Our multi-sample platform demonstrates effective upgrades to conventional single-sample imaging platforms and paves the way for diverse quantitative long-term imaging studies.

**Summary statement:** We present a dedicated multi-sample light sheet imaging, processing and analysis platform and demonstrate its value for studies of vascular growth in zebrafish.

## Introduction

The cardiovascular system is among the earliest functional organs to be formed during vertebrate development. From a few individual mesodermal precursor cells, a complex network of vessels forms through a variety of morphogenetic processes (Ellertsdottir et al., 2010; Gore et al., 2012). While many molecular actors involved in its formation have been identified (Adams and Alitalo, 2007; Gore et al., 2012; Hogan and Schulte-Merker, 2017), much remains unknown about how this intricate network comes into shape on the whole embryo level. A particularly exciting open question is how the volume of the vascular system changes over development. Vascular volume is a strong proxy for overall vascular system size and consequently its development provides insight into fundamental aspects of tissue development and whole embryo morphogenesis. Furthermore, as the vasculature is a closed system, its volume is closely linked to blood pressure and flow characteristics. Changes in volume might therefore be involved in regulation of crucial steps of vascular formation. Here, we show how a combination of dedicated multi-sample preparation, comprehensive imaging and data processing, a novel segmentation approach, and growth data analysis provides a precise and quantitative characterization of embryonic vascular volume development.

The zebrafish has become an especially valuable tool to understand vascular formation on the whole embryo level, with many available vascular transgenic lines (Chávez et al., 2016). Compared to other vertebrate model organisms such as mice, the development of zebrafish is fast, outside of the mother, and the optical translucency of zebrafish embryos provides an ideal setting for long-term in-vivo time-lapse imaging experiments (Kimmel et al., 1995).

For long-term live imaging of zebrafish embryos and larvae, light sheet microscopy (Huisken et al., 2004; Keller, 2013) has become the method of choice due to its illumination and detection scheme that provides minimal photo-bleaching and phototoxicity (Daetwyler and Huisken, 2016; Icha et al., 2017; Power and Huisken, 2017). Moreover, light sheet microscopy offers sample rotation in the microscope for multi-view imaging and tiling for full embryo coverage (Weber and Huisken, 2012), which is a necessity for imaging the entire vascular system. In addition, imaging zebrafish embryos over several days requires a sample embedding technique that provides mechanical constraints to ensure proper sample orientation but does not limit oxygen access or restrict growth. For single samples, an embedding method using fluorinated polypropylene (FEP) tubes (Kaufmann et al., 2012) has been widely accepted. However, to quantitatively analyze the observed processes, more than one sample needs to be imaged. Ideally, several samples are imaged in one experiment simultaneously, which is especially important and efficient if the experiment takes several days to complete. Therefore, multi-sample imaging is highly desired for long-term imaging studies.

The number of samples during imaging can be increased either by delivering samples with flow (Gualda et al., 2015; Regmi et al., 2013; Regmi et al., 2014; Wu et al., 2013) or by embedding multiple samples at the same time (de Luis Balaguer et al., 2016; Schmid et al., 2013). Delivering samples by flow does not offer the precise control of sample orientation needed for optimal image quality and comparison of different time points. Therefore, multi-sample embedding solutions are more promising for long-term imaging studies. However, existing multi-sample embedding solutions are currently only suitable for embryos still in their protective envelope, the chorion, (Schmid et al., 2013) but not for growing zebrafish larvae. Sample holders for several plants have been designed to allow plant growth in near physiological condition (de Luis Balaguer et al., 2016) but do not provide sample rotation during imaging to access the entire samples. Therefore, the challenge is to develop a multi-sample embedding technique that allows for multi-view imaging.

A further complication arises when imaging several embryos simultaneously as data handling becomes more challenging, with datasets easily exceeding several terabytes in size. Furthermore, such datasets are comprised of many acquisition volumes, angles, and time points over multiple samples. It is consequently not possible to load an entire experiment into computer memory to inspect the data and apply conventional data processing and visualization workflows. Therefore, custom data processing tools are needed to automatically generate 3D stitched datasets for further analysis, and the visualization thereof for easy inspection of acquired data.

Next, for an accurate description of vascular volume changes, visual inspections and qualitative analysis are not sufficient. Quantitative measurement of the vascular volume requires segmentation. Most segmentation approaches in developmental biology focus on analyzing nuclei and cytoplasmic content (Amat et al., 2014), but segmentation of the vasculature is more challenging due to the variety of vessel sizes, intensity changes over vessel walls and, most importantly, due to the hollow-tube structure of vessels. Therefore, no reliable segmentation approach of endothelial signal over long time periods has yet been established. To help with segmentation, micro-angiography (Isogai et al., 2001) is often used, in which a fluorescent dye is injected into the vasculature. However, at early developmental stages, the vasculature is not completely closed and dye rapidly leaks into the surrounding tissue, rendering microangiography not applicable for long-term imaging studies. A novel strategy of vascular segmentation is thus required to extract quantitative growth measurements of the whole vasculature.

To interpret the quantitative measurements resulting from the complete multi-sample imaging and processing platform, there are many established mathematical growth models available (Hernandez-Llamas and Ratkowsky, 2004; Tsoularis and Wallace, 2002). They broadly fall into two groups, namely deterministic models originating from integration of a (partial) differential growth equation and stochastic models. However, none of these models were capable of sufficiently capturing or describing our data. We therefore suggest a new class of stochastic growth models based on logarithmic rescaling of time to describe embryonic vascular development.

## RESULTS

### Multi sample preparation for long-term time-lapse imaging

Successful vascular imaging relies on immobilization of the sample and mechanically constraining it to the field of view through embedding. In contrast to the widely established zebrafish embedding protocol (Kaufmann et al., 2012), we abstained from using tricaine. Instead, to immobilize the embryos, we injected them with α-bungarotoxin RNA (Swinburne et al., 2015) at the one cell stage. To embed several samples for simultaneous long-term imaging, we adapted the established protocol of single embryo embedding. We first embedded individual zebrafish embryos in 0.1% agarose inside of FEP tubes for long-term imaging as described (Kaufmann et al., 2012) (Fig. 1A). These FEP tubes were cut to a length of about 8 mm and several of them were attached with other larger FEP tubes as connectors (Fig. 1B). Holes in these connectors ensured exchange of oxygen and liquids during imaging. A tight fit was ensured by using FEP’s elastic properties that allow specification of the inner diameter of the connector slightly smaller than that of the outer diameter of the tube containing the fish. This embedding technique was readily available, flexible, and provided stable embedding of up to five fish for long-term imaging in one tube assembly with a length of about 6 cm. A detailed step-by-step protocol for multi-sample embedding can be found in the supplementary material (S1).

**Figure 1:**
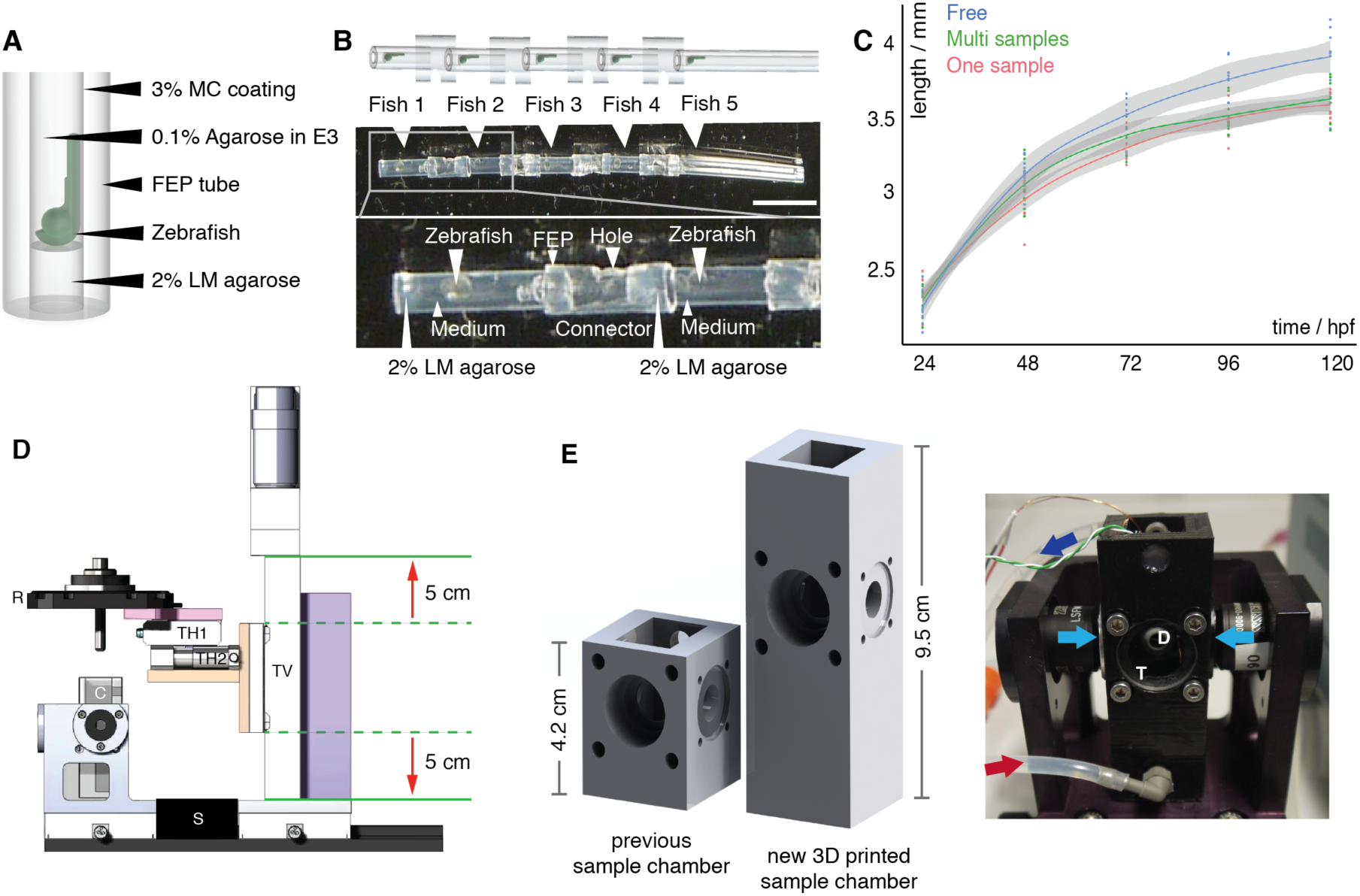
Multi-sample embedding and necessary microscope modifications. (**A**) Schematic of embedding of one zebrafish embryo in fluorinated polypropylene (FEP) tube using 2% low melting (LM) agarose as plug, 0.1% agarose in E3 as medium and 3% methylcellulose (MC) as coating of the FEP tube as described by (Kaufmann et al., 2012). (**B**) Schematic (top) and picture (middle) of 5 mounted zebrafish (white arrowheads) in FEP tube pieces assembled with FEP connectors. The enlarged region (bottom) shows details of the embedding: zebrafish mounted in the FEP tubes rested on a 2% LM agarose plug and were embedded in E3 medium containing 0.1% agarose. FEP tubes containing a single zebrafish embryo were attached with FEP connectors containing a hole. Scale bar: 1 cm. (**C**) Growth curve of freely swimming fish (blue, n=8), single embedded samples (red, n=10), and samples embedded in the multi-sample tube (green, n=9) with the 0.95 confidence interval of Loess interpolation in gray. (**D**) The new translational stage design of the multi-sample imaging platform ensured a vertical travel range of 10 cm (red arrows) and was built with custom parts, e.g. adapter of the rotational stage [R] onto the translational stage platform (pink), adapter of horizontal translational stages [TH1, TH2] onto the vertical translational stage (TV, orange), and a stage mount (purple). Spacer S, sample chamber C. (**E**) Comparison of a traditional one-sample SPIM chamber with the new 3D-printed sample chamber and a picture of the new sample chamber connected to a perfusion system with inflow (red arrow) and outflow (dark blue arrow), the two illumination objectives for dual-sided illumination (light blue arrows), a window for transmission (T) and a detection objective (D) in the back.

To ensure that the multi-sample embedding did not compromise growth, we measured the overall body length of freely swimming zebrafish, single-sample and multiple-sample preparations over time and compared their growth curves (Fig. 1C). Until 24 hours after embedding (48 hours post fertilization [hpf]), no growth difference was detected (ANOVA analysis, p-value 0.1721). This changed at 72 hpf (ANOVA analysis, p-value 0.0013), when the embedded fish were 5% smaller after 2 days of embedding than the freely swimming fish. However, the data provided no evidence that there was a growth impairment of the multi-sample embedded fish compared to the established one-sample embedding technique (two-sided t-test between one-sample and multi-sample embedding: p-value 0.06 at 72 hpf, 0.93 at 96 hpf, and 0.98 at 120 hpf). Moreover, no additional growth defects such as edemas were detected by visual inspection (Fig. S4).

### Hardware adaptations for multi-sample imaging

The new 6 cm long tube assembly containing the embedded zebrafish embryos did not fit on our custom and other commercially available light sheet implementations. Therefore, we decided to upgrade an existing custom three-lens mSPIM (Huisken and Stainier, 2007) that has been successfully applied in long-term imaging studies (Lenard et al., 2015). The existing mSPIM was equipped with two illumination and one orthogonal detection arms. Furthermore, its rotational stage enabled sample rotation so that zebrafish embryos could be oriented to image them for ideal penetration and coverage. To upgrade this custom system, we designed a new translational stage system (Fig. 1D) with larger travel ranges to move every one of the five samples into the microscope’s field of view. Furthermore, the depth and height of the sample chamber (Fig. 1E) was increased to provide enough space for the tube assembly. The taller chamber was 3D-printed (Fig. 1E), making the chamber a cost-effective and easily adaptable unit. We incorporated in- and outlets for a perfusion system into the sample chamber to allow for temperature control of medium and sample.

### Multi-sample imaging of several zebrafish embryos

To study the development of the vascular volume, we simultaneously imaged five zebrafish embryos expressing a green fluorescent vascular endothelial marker, *Tg(kdrl:EGFP)* (Jin et al., 2005), and a red blood cell marker, *Tg(GATA1a:dsRed)* (Traver et al., 2003). The green fluorescent marker labelled the vessel walls and thus the outlines of the vascular system. The red blood cell marker labelled erythrocytes and thus provided information about the inside of the vasculature.

To achieve optimal coverage of the entire vascular system, we chose three optimal angles for imaging (Fig. S5): To capture a good view on the head vasculature, we imaged the embryos with dual-sided illumination and the detection oriented dorsally. To achieve a good whole-embryo view, we imaged the zebrafish from two opposite angles ±60 degrees rotated from the sagittal plane with single-sided illumination.

The EMCCD acquisition cameras only had a chip size of 960×960 pixels corresponding to 1.097×1.097 mm field of view. As the embryos grew to a size of about 3.5 to 4 mm at 5 dpf, several acquisition volumes were required to reconstruct the whole embryos. Together with an overlap of about 30% for reliable data stitching, five acquisition volumes were needed to image one whole embryo from one angle. Therefore, for imaging five embryos from three angles, 75 acquisition volumes were required. To capture the total 3D volume, we imaged every acquisition volume with at least 200 z-planes separated by 4 μm.

To image the development of the vascular volume over time, we began imaging at around 17 hpf when the cells started to fuse to larger vessels by in-situ aggregation of angioblasts (Ellertsdottir et al., 2010). To understand the overall vascular volume changes rather than obtaining a movie of the behavior of one individual vessel, we imaged the entire developing vasculature every 20 min over a time period of at least three days. With these settings (75 acquisition volumes, dual-color, 200 planes/acquisition volume, time step of 20 min), the corresponding data generation rate amounted to approximately 3.5 TB / day.

### Microscope software adaptations for multi-sample imaging

The high data rate from our multi-sample imaging SPIM instrument posed a considerable challenge as hard-drives available at the microscope had a total capacity of only 6 TB, making a three-day timelapse experiment impossible. We therefore decided to copy the accumulated data from every time point to a large, central storage in-between acquisitions (material and methods, Fig. S6). This approach also eliminated the usual time-consuming data transfer at the end of an experiment, which prevents the immediate start of the following experiment. Copying data during an experiment thus also increased the overall throughput of the microscope.

We further implemented an automated mosaic generation tool as manually configuring a high number of individual acquisition volumes for each experiment is very time consuming and error prone. Given a starting position, e.g. the head of the fish, the other acquisition volume’s positions were automatically determined with a 30% overlap, ensuring reliable stitching during data processing. The overall number of acquisition volumes was selected based on the expected growth of the zebrafish over the course of the time-lapse at the start of the experiment.

### Dedicated processing pipeline

Multi-sample acquisition over three days as described above resulted in large datasets of over 10 TB with a convoluted data structure comprising of many acquisition volumes, several angles and fish, and two channels. Using custom modular ImageJ / Fiji plugins (Schindelin et al., 2012), we transformed this imagery dataset into a dataset that contained one stitched 3D stack per timepoint, channel, angle and fish (Fig. S7). Furthermore, to visually check the data quality, we generated maximum intensity projections that also enabled a qualitative description of the formation of the vasculature (Movie 1, Movie 2). We adapted the plugins for use on a high-performance cluster for fast processing (S5). Using the microscope stage parameters, we also automated all processing steps so that only the folder of the experiment had to be indicated for processing and visualization. The code for all processing plugins is freely available at: https://github.com/DaetwylerStephan/multi_sample_SPIM.

### Segmentation of the vascular data

To quantitatively measure the vascular volume changes over time, a segmentation of the vasculature was required. Vessel segmentation is challenging due to vessel geometry and characteristics of the fluorescent marker (Fig. 2A,E). Blood vessels are hollow tubes of widely varying diameters, often in very close proximity to each other. Additionally, the endothelial marker showed heterogeneous densities across the vessel walls, e.g. with higher intensities around nuclei. Therefore, classical approaches such as filter-based approaches (Sato et al., 1998) or simple thresholding did not result in good segmentations (data not shown).

**Figure 2:**
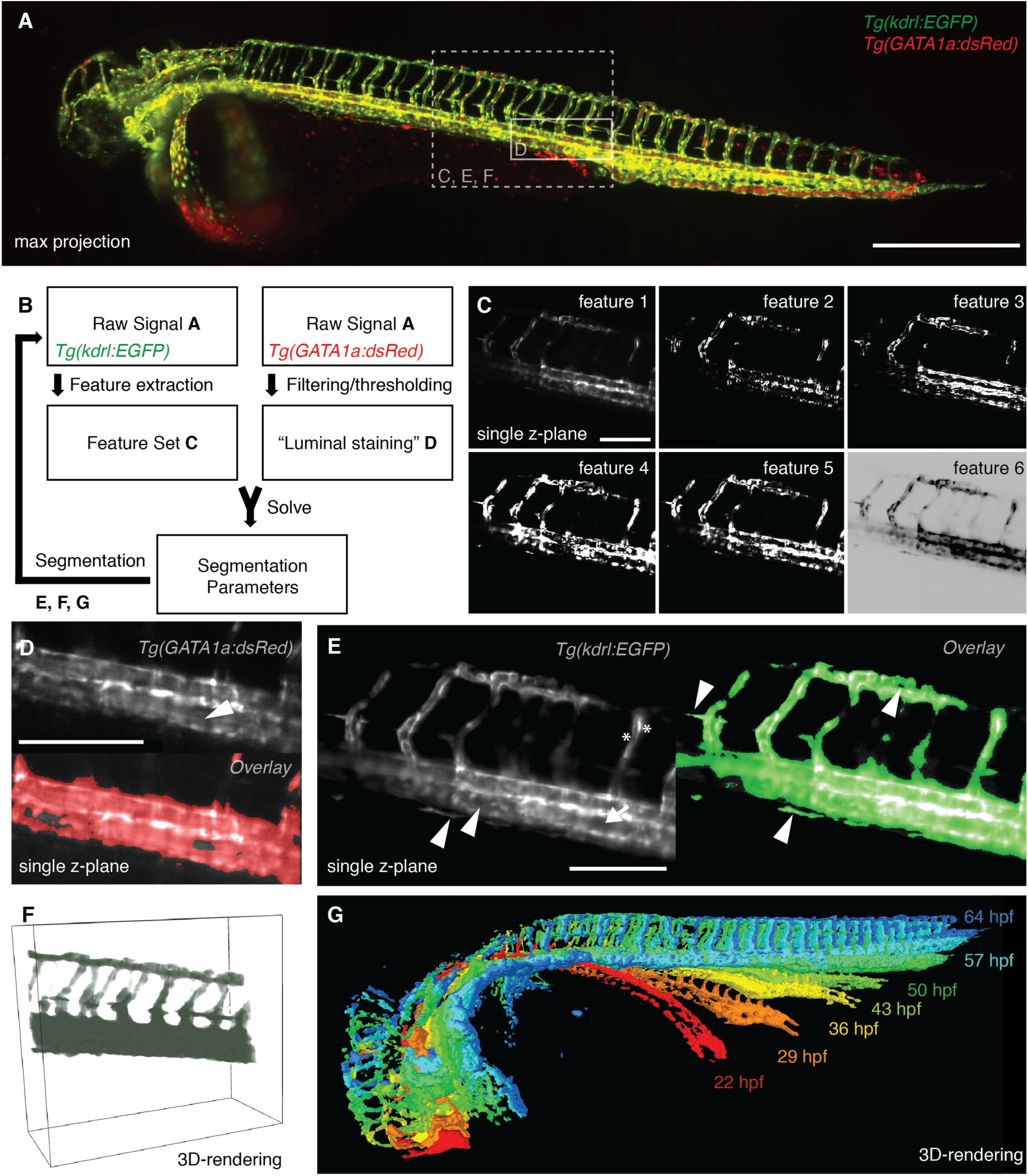
Segmentation of the vascular data. (**A**) Maximum intensity projection of a zebrafish expressing the endothelial marker *Tg(kdrl:EGFP)* in green and the red blood cell marker *Tg(GATA1a:dsRED)* in red with boxes depicting selected regions shown in (C-F). Scale bar: 500 μm. (**B**) Schematic of the segmentation process. (**C**) Features extracted from the endothelial marker *Tg(kdrl:EGFP)*, with the raw signal (feature 1), the gradient x (feature 2), gradient y (feature 3), gradient z (feature 4), total gradient (feature 5) and the inverse gradient weighted raw image (feature 6). Scale bar: 150 μm. (**D**) Single plane of a 3D stack of the red blood cell marker *Tg(GATA1a:dsRED)* (top). As the red blood cells circulate in the vasculature, the interior of vessels was also filled with fluorescent signal (white arrowhead). Therefore, filtering and thresholding of the red blood cell marker raw signal (top) provided a ground truth (bottom, red) of signal inside the vasculature from which the segmentation parameters could be calculated. Scale bar: 150 μm. (**E**) Single plane of a 3D stack of the endothelial marker (left) highlighting the challenges of vascular segmentation: hollow tubes (arrow), intensity differences (asterisk), small vessels next to large vessel (arrow head). With our segmentation approach (right), even fine structures of the vasculature were segmented correctly (arrowheads). Scale bar: 150 μm. (**F**) 3D-rendering of the selected region with the segmentation in green color. (**G**) 3D-rendering of the segmentations at different time points over development.

To provide an effective segmentation, we designed a novel approach by complementing the signal of the endothelial marker with the signal from the red blood cells *Tg(gata1a:DsRed).* Red blood cells are inherent proxies for luminal markers as they circulate inside the vasculature. Consequently, we used a segmentation of the red blood cells as training data for a machine-learning based approach of vessel segmentation on the endothelial signal. Segmentations were performed by computing a set of feature images based upon gradients of the *Tg(kdrl:EGFP)* channel, and solving for weights of the feature images that predicted the locations of red blood cells based on the *Tg(GATA1a:dsRed)* channel. The calculated weights were then applied on the whole 3D stack to obtain a segmentation of the vascular volume (see methods for details).

We implemented our efficient and fully-automated image segmentation pipeline in FunImageJ (Harrington et al., 2018). FunImageJ was used with a standard distribution of Fiji (Schindelin et al., 2012), and as a standalone program on a high-performance computing cluster to enable parallel processing of whole datasets.

### Vascular growth curve rates

With the segmentation in hand (Fig. 3A), quantifying the overall growth of the vascular system was a straightforward process of counting the number of segmented voxels in isotropic 3D stacks. To temporally align the volume measurements of different fish, we selected the anastomosis of the left and right primordial hindbrain channel (PHBC) as reference points.

**Figure 3:**
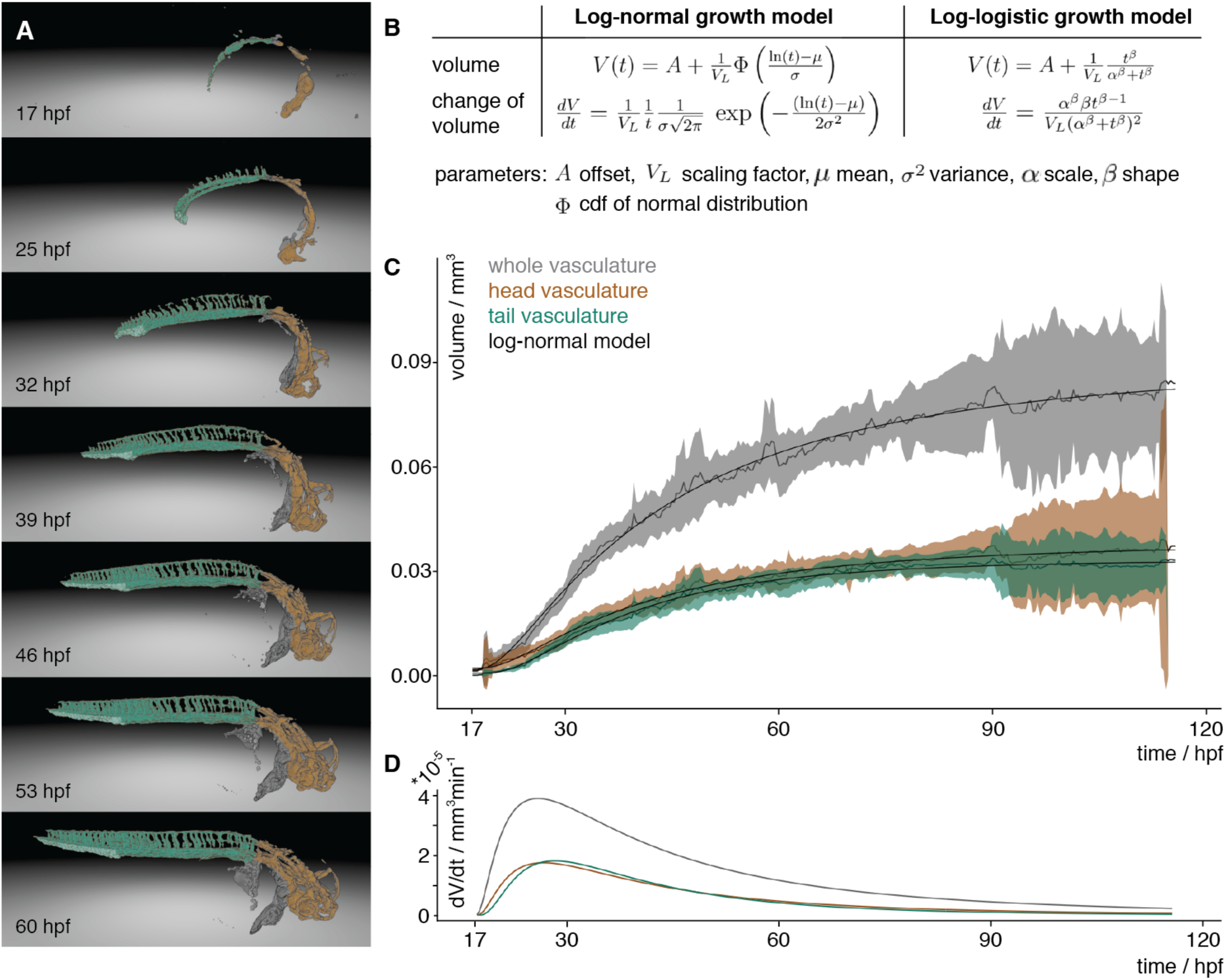
Vascular volume growth characteristics of zebrafish. (**A**) Segmentation of the vasculature at seven different time points labelled with the annotation of the head (orange) and tail (turquoise) with its caudalveinplexus (light-turquoise) and rest (grey). (**B**) Equations of the cumulative log-logistic and log-normal growth models describing the volume V(t) and change of volume dV/dt over time t with offset A, scaling factor V_L_, mean and standard deviation of log-normal distribution μ, σ and Φ cumulative distribution function (cdf) of the standard normal distribution, and log-logistic scale parameter α and shape parameter β. (**C**) Experimental measurements of the volume over time of the whole vasculature (gray), the head (brown) and tail (turquoise) vasculature. The mean of the measurements is depicted with a solid line and the 95% confidence interval (t-statistics, n=7) as a ribbon in the corresponding color. The black line depicts the approximation of the volume by the log-normal growth model. The same panel for the log-logistic model is in the supplementary (Fig. S10A). (**D**) The volume growth rate of the whole vasculature (gray), head (brown) and tail (turquoise) was calculated by inserting the parameters obtained from the approximation into the change of volume equation of the log-normal growth model. The same panel for the log-logistic model is in the supplementary (Fig. S10B).

The resulting quantitative data revealed that the growth of the vascular volume was well described by a group of models relying on logarithmic rescaling of time including the cumulative log-logistic and the cumulative log-normal growth model (Fig. 3B). To account for a limited time window of observation, a scaling parameter and an offset were introduced in both models. The offset *A* described the lower asymptote i.e. the vascular volume already formed at the start of an imaging experiment. The scaling factor *V*_*L*_ described the upper asymptote i.e. the maximum volume reached during the observed development. For the log-normal growth model, two parameters μ (mean) and σ (standard deviation), which define the cumulative log-normal distribution function, were required to describe volume growth (Fig. 3C). For the log-logistic model, the scale parameter α and the shape parameter β were required (Fig. S10). Consequently, in total only 4 parameters were required to model the development of the vascular volume over time. We used non-linear minimization to obtain the fitting parameters for both models (Table 1).

**Table 1.**
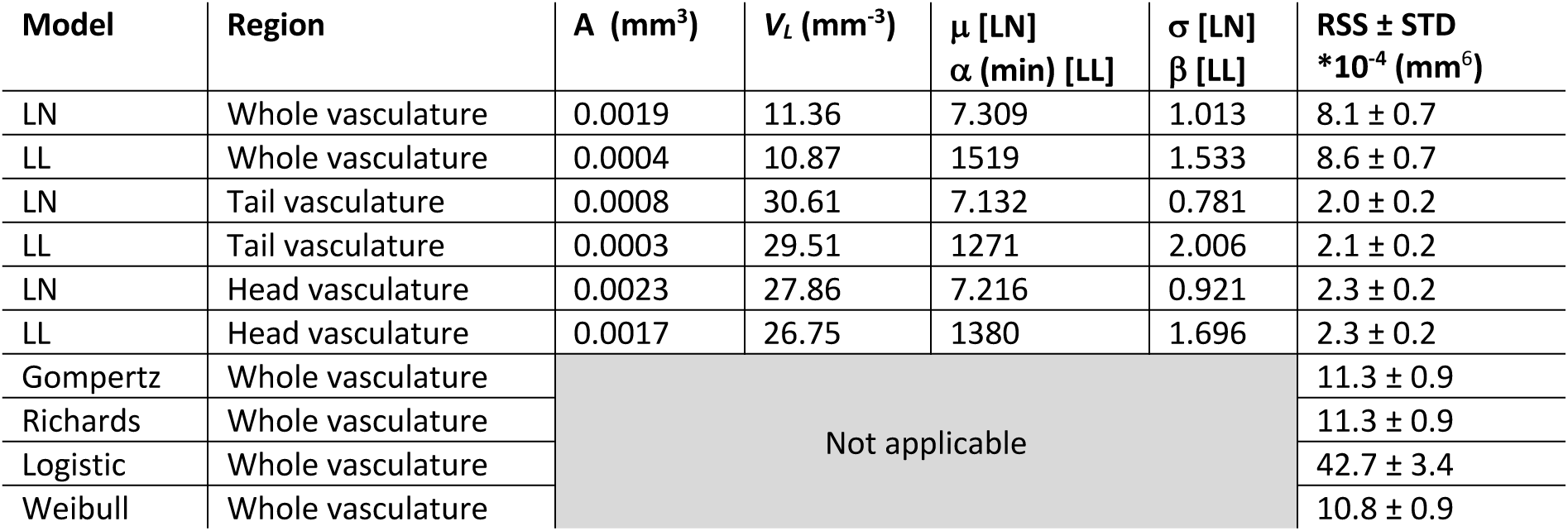
Fitting parameters for volume growth of log-normal (LN) and log-logistic (LL) growth models

We compared the cumulative log-normal and cumulative log-logistic growth model with other established growth models by calculating the residual sum of squares (RSS). A small RSS indicates a good fit to the data. The comparison of the RSS values revealed that the cumulative log-normal growth model and the log-logistic model of the whole vasculature volume growth exceeded other established growth models such as the Gompertz Model, logistic growth, Janoscheck and Weibull Model, or the Richard’s model (Table 1, Fig. S9). Moreover, in comparison to these models, visual inspection of the residuals of the cumulative log-normal and growth model indicated that the log-normal model indeed fit best (Fig. S9).

To understand whether the overall growth characteristics of the whole vasculature are reflected in the growth of its subnetworks such as the head or tail vasculature, we established a manually-curated annotation of the vasculature (Movie 3). The volume changes of the differently annotated regions were determined by counting the segmented voxels with the corresponding annotation label in isotropic 3D stacks. The analysis of vascular volume development of the head and tail vasculature (Fig. 3C) revealed that they were also well described by the cumulative log-logistic and log-normal models. The fitting parameters for both models were obtained by linear minimization and revealed a good fit (Table 1).

We inserted the above obtained parameters into the scaled log-normal function on which the cumulative log-normal model was based (Fig. 3B). The scaled log-normal function was the derivative of the cumulative log-normal model and thereby revealed the growth rate of the vascular volume (Fig. 3D). The maximal growth rate was obtained to be around 26 hpf for the whole vasculature, 27 hpf for the head and 28 hpf for the tail vasculature. Before this, the growth rate rapidly increased while after reaching the maximum, the growth rate slowly decreased. Inserting the parameters of the cumulative log-logistic model into its derivative revealed the same values for whole vasculature and head vasculature, and 29 hpf for the tail vasculature.

## DISCUSSION

We have presented a dedicated and complete workflow of multi-sample preparation, multi-sample imaging, data processing and quantitative analysis of the vascular volume. Several key innovations in all those disciplines were necessary: We introduced a new multi-sample embedding protocol, an upgraded light sheet microscope, a comprehensive library of data processing plugins, a novel vascular segmentation approach using inherent biological markers and new growth models relying on logarithmic rescaling of time for describing the development of embryonic vascular volume. The presented tools will further open the door for many other long-term imaging studies in fields such as developmental biology or xenograft models in cancer biology.

### Growth of the vasculature

With our multi-sample imaging and analysis platform, we obtained a high-quality data set describing the growth of the vascular volume over time. Descriptive features of the observed time course include a slow growth at the beginning, a maximum volume growth rate at around 26-29 hpf and then a decline of the growth rate resulting in a saturating growth process. To describe this growth, we fit well-established growth models (Fig. S9) that have been used for saturating growth processes such as the logistic function (Table 1) and found that they did not reproduce the first hours of vascular development well. In searching for an alternative growth model, we found that the best growth models to explain the data all include an effective, logarithmic rescaling of time. Log-normal-like dynamics and log-logistic growth fit best and are clearly distinguishable from other, non-logarithmic growth models, but not from one another. To our knowledge, this is the first time that such models have been shown to be applicable to biological growth data.

Logarithmic behavior often comes into play when proportional effects are driving the underlying processes, i.e. when a quantity X_t_ at time *t* is a proportion e_t_ of its size X_t-1_ at a previous time *t*-1 such that X_t_ = X_t-1_ + e_t_ X_t-1_ (Graham et al., 2003). In vasculature development, we speculate that factors such as (i) nutrient availability or (ii) gene expression can introduce proportionality and thereby explain the observed logarithmic rescaling behavior. (i) A limited store of nutrients is deposited by the mother and remains the predominant source of nutrients over the first days of development. Assuming an exponential usage of nutrients out of this store, the available remaining nutrients are proportional to the original amount deposited as development progresses. In such a scenario, the growth steps over time would be linked to the decay and thereby become logarithmically rescaled. (ii) Also gene expression could introduce similar proportionality effects. Here, feedback loops in gene expression patterns would enforce dependence on the pattern at a previous time point.

### Blood flow and vascular volume development

Interestingly, the maximum volume growth rate at around 26-29 hpf corresponded to the onset of blood flow in the whole network. Blood flow might therefore be involved in a global change of vascular growth characteristics such as gene expression patterns. Indeed, it is well known that endothelial cells can sense blood flow (Franco et al., 2016), and that blood flow plays a role in vascular plexus remodeling (Lenard et al., 2015). On a global level, flow might induce two regimes of vascular growth: a first phase independent of flow with high growth rate and a second phase with reduced growth rates where flow plays a crucial role. To further investigate flow dependence of the network, our quantitative volume measurements will pave the way for new accurate data-driven computer simulations as they provide complete vessel boundaries and volume measurements for the whole embryo. This will enable simulations of shear stresses and blood velocity over the whole vasculature given a certain heartrate, and consequently reveal correlations between shear stresses and vascular remodeling. To validate the simulations, the already acquired red blood cell data could be used to extract blood flow velocities at different parts of the vasculature and compare it to the simulation’s predictions.

### Growth of vascular subnetworks

The volume measurements of large subnetworks of the vasculature such as the head and the tail vasculature were also well described by a cumulative log-normal function. This indicates that also those large subnetworks were subject to the same growth constraints as the whole vasculature and explain in parts the fractal nature of vasculature. However, these constraints might not apply to smaller subnetworks as they can draw from and release resources to neighboring parts of the network, e.g. by cell migration. Therefore, those smaller subnetworks might show different growth behaviors. Indeed, the caudalveinplexus located in the tail of the vasculature first expanded, then remodeled and after 73 hpf decreased in size (Fig. S11) and therefore did not follow a log-normal growth rate.

### Multi-sample capacity is desired and important

Simultaneous imaging of several samples offers higher efficiency, shorter overall experimental time and thus lower costs than sequential image acquisition. Furthermore, it enables the study of embryos from the same parents, growing in the exact same conditions. This is especially important for imaging mutant fish lines. Homozygous mutant fish are often not viable (Kim et al., 2011), and therefore heterozygous fish lines are grown. However, only one quarter of their progeny are homozygous for the mutation of interest and exhibit the phenotype. To observe its establishment, fish are ideally selected before a phenotype is observable. Consequently, multi-sample imaging is essential for studying mutant fish lines with high spatial and temporal resolution. Moreover, with the multi-sample platform, new studies will be possible to understand variation and robustness of a biological structures. Indeed, the vasculature of zebrafish also shows variation (Fig. S12). Our long-term time-lapse platform will reveal how such patterns of variation are established, maintained and/or remodeled.

The ability to image many samples is also important for experiments that are time consuming in preparation and/or rely on the short-term availability of rare, restricted, or otherwise difficult to obtain samples. Such samples include patient-derived samples, xenotransplantation studies, observations of induced cancer cells or small-scale screens of selected compounds. Our multi-sample platform will provide the required sample capacity to tackle such projects.

### Integration is critical

The power of our multi-sample imaging platform lies in the integration of all of the important steps of multi-sample imaging: sample preparations, imaging, data processing and data analysis into a single dedicated pipeline to obtain quantitative data (Fig. 4). For example, the data analysis benefits from the right choice of sample. Only by using the red blood cells as a luminal marker, we obtained a good training data to solve for the segmentation parameters. In a further example, imaging and data processing are interlinked as the acquisition parameters, here the translational stage positions, are crucial for reliable stitching. As all of our processing and analysis code is freely available, other researchers might adapt parts of the whole pipeline to their dedicated research question. Moreover, our suggested upgrade for multi-sample imaging might be easily incorporated into other custom-built systems, together with the here in-detail described embedding protocol. Hence, this toolset will help current imaging strategies move from qualitative descriptions of single observations to quantitative analysis over multiple samples.

**Figure 4:**
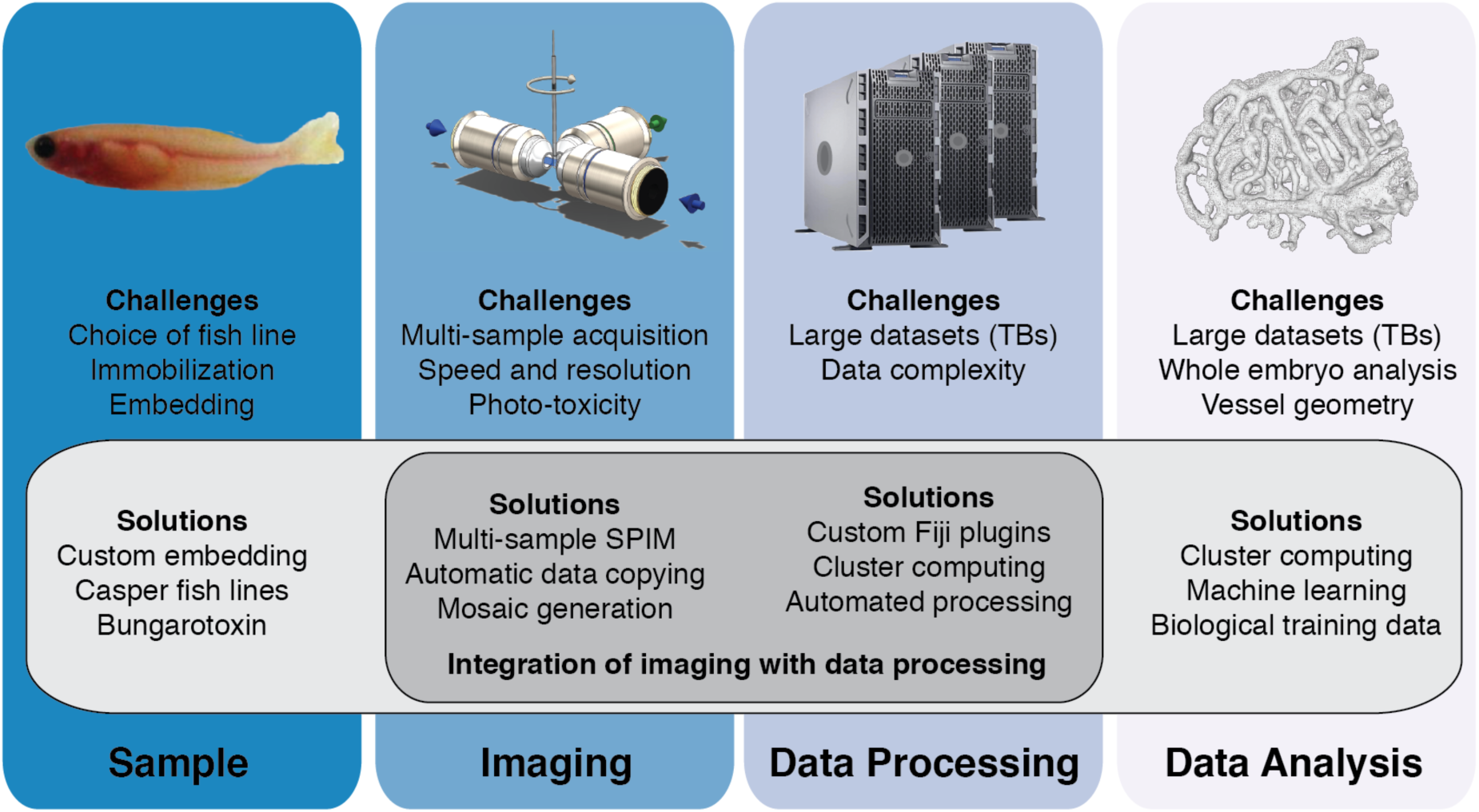
A dedicated imaging, processing and analysis platform for multi-sample imaging. Integration of important steps of multi-sample imaging such as sample preparation, imaging, data processing or data analysis facilitates the individual steps and enables multi-sample imaging.

## MATERIALS AND METHODS

### Ethics Statement

The animal experiments were performed in accordance with the European Union (EU) directive 2011/63/EU as well as the German Animal Welfare Act.

### Zebrafish sample preparation

Zebrafish (*Danio rerio*) adults and embryos were kept at 28.5 °C and were handled according to established protocols (Nüsslein-Volhard and Dahm, 2002; Westerfield, 2000).

To understand vascular growth in zebrafish, we compared *Tg(fli1a:EGFP)* (Lawson and Weinstein, 2002) and *Tg(kdrl:EGFP)* (Jin et al., 2005). As the marker *Tg(fli1a:EGFP)* was expressed not only in the head vasculature but more broadly in the head (data not shown), we decided to use *Tg(kdrl:EGFP)* and crossed this fish line into a casper (White et al., 2008) background to suppress formation of pigmentation. The *Tg(kdrl:EGFP)* casper fish line was crossed with the line *Tg(GATA1a:dsRed)* (Traver et al., 2003) expressing a fluorescent red blood cell marker. For time-lapse experiments, zebrafish embryos were injected at the one-cells stage with 30 pg of α-bungarotoxin RNA (Swinburne et al., 2015) to ensure immobilization during the time-lapse.

A detailed step-by-step protocol for embedding the embryos for imaging is in the supplementary (S1).

### Growth measurement

Freely swimming zebrafish, single-sample and multiple-sample preparations were set up at 24 hpf. Freely swimming zebrafish (n=8), single-sample (n=10) and multi-sample (n=9) preparations were then imaged using an AVT stingray camera connected to an Olympus SZX16 stereo microscope at 24, 48, 72, 96 and 120 hpf. To calibrate the length measurement, a PYSER-SGI stage micrometer (10 mm/0.1 mm) was imaged together with the zebrafish embryo. Fiji (Schindelin et al., 2012) was used to first calibrate and then measure the length of the zebrafish embryo.

### *In vivo* Time-Lapse Imaging

Long-term time-lapse imaging was performed on a home-built multidirectional SPIM (mSPIM) setup (Huisken and Stainier, 2007) upgraded to multi-sample capacity (result section). The microscope was equipped with two Zeiss 10×/0.2 NA illumination objectives and an UMPlanFL N 10×/0.3 NA Olympus detection objective. Two Coherent Sapphire 488 nm −100 CW / 561 nm – 100 CW lasers were used to illuminate the sample. The images were recorded with two Andor DV885 iXon EM-CCD cameras. The embryos were imaged at least every 25 min for up to 4 days starting around 17 hpf.

### Design of the translational stage system

The translational stage system of the microscope required high precision as it positioned the sample and scanned it through the light sheet. Therefore, a precise, translational stage with a longer travel range had to replace the existing vertical translational stage unit (Fig. 1D). We chose the M-404.4PD precision translation stage (Physik Instrumente) as it offered a unidirectional repeatability of 0.5 μm equal to half a pixel on the camera, and an overall travel range of 100 mm, which was sufficient for imaging several samples. To integrate the larger stage, we designed custom parts to connect the M-404.4PD stage to two M-111.1DG translational stages (Physik Instrumente) for lateral and axial scanning. We further added a solid metal block to stabilize the translational stage system and avoid vibrations caused by translational movements.

### Data copying in-between timepoints

As copy tool we used the Windows command line executable Robocopy that was started right after the acquisition of every 3D stack and copied data until just before a new stack acquisition was started. Robocopy was integrated into Labview, the microscope control software, by their executable interface framework. This ensured robust data transfer as Robocopy only removed old data once the integrity of the file at the new server location was checked.

### Data processing

Custom-made data processing tools in Fiji (Schindelin et al., 2012) were written to process the data from the microscope automatically. For visualization, the acquired fluorescence images were projected using maximum intensity projections. The resulting projections were stitched and fused with linear blending using custom code adapted from the Fiji stitching plugin by Stephan Preibisch et al. (Preibisch et al., 2009) that relies on phase correlation (Kuglin and Hines, 1975). For successful stitching (Fig. S8), we initialized the stitching with the translational stage positions. As the translational stages were very precise, the stitching was robust and determined globally for all timepoints. The stitching parameters of the maximum intensity projections were also applied to the 3D data to generate one 3D fused stack per timepoint, angle, fish and channel. While the different channels were already aligned optically, we used a manual GUI interface to obtain a precise fine alignment of the different channels using rigid registration. To reduce the amount of data for storage to about half the size, the data was compressed by zipping.

Code of the custom-made processing steps is available freely at: https://github.com/DaetwylerStephan/multi_sample_SPIM

### Segmentation

The image segmentation software was developed using FunImageJ (Harrington et al., 2018), a Lisp-based interface for ImageJ. Code is freely available at: https://github.com/kephale/virtualfish-segmentation

The segmentation parameters were determined for each individual 3D stack by applying a machine-based learning approach. Candidate locations for sampling training data were selected by taking the conjunction of the thresholded red blood cell and vasculature channels. Sample points were chosen by randomly sampling candidate coordinates and collecting the first N positive points and first N negative points without replacement (N=5,000).

The sample points were used as a target for the parameter fitting procedure. Feature vectors were computed for each sample point using first-order moments calculated with ImgLib2 (Pietzsch et al., 2012). The features encoded the original image, X, Y, and Z gradients of the vasculature channel, the total gradient magnitude, and an inverse-gradient weighted version of the original image. The feature vectors for all sample points were then composed as a matrix and a target vector was generated, representing the positive/negative labels of training points as 1 and 0, respectively. Singular value decomposition was then used to solve for a set of feature weights that maximized predictions of the target vector value.

The vector of feature weights obtained by solving the linear fitting procedure was then used to compute an image of segmentation scores. Images were segmented by first using the fitted feature weights to compute a linear combination of the feature maps. The resulting image encoded the segmentation score for each voxel. The optimization procedure did not guarantee that the resulting outputs were restricted to [0,1], therefore it was not a probability map. The segmentation score image was then thresholded using the Triangle algorithm (Zack et al., 1977) resulting in a binary labeling of the image. A morphological erosion followed by a dilation filtered out single-pixel fragments. Meshes were generated for visualization from binary segmentations using the marching cubes algorithm from ImageJ2 (Rueden et al., 2017).

### Visualization of segmentation

To visually check the quality of the segmentation (Fig. 2E), we overlaid the segmented images with the raw signal at selected timepoints. We further visually inspected all segmentations over the whole time-lapse course by maximum intensity projections of them, and by 3D rendering (Fig. 2F,G, Movie 3) the segmentation and the annotations using SciView (available at https://github.com/scenerygraphics/sciview).

### Quantification of vascular growth

For quantification of the segmentation, we rescaled each segmented 3D stack of a time-lapse series to isotropic resolution and then counted the number of segmented voxels. We quantified only the two opposite angles rotated by ±60 degrees from the sagittal plane. Those two angles provided the best resolution of the whole embryo zebrafish vasculature. To obtain the vascular volume growth curves for one fish, the quantification result of both angles was averaged.

### Annotation of the segmented vasculature

For the annotation, we used the maximum intensity projections of the endothelial signal over time. We first selected manually a region of interest such as the caudalveinplexus, the head or the tail vasculature (including the caudalveinplexus) at the last timepoint of a time series. To automatically track this region of interest over time, we sequentially determined the region of interest at time t-1 given the region of interest at time t. For this, the boundary of the region of interest was discretized by points and for each point of the boundary the corresponding point at time t-1 was determined. Assuming that only small-scale changes were present, the computation for each point was reduced by only considering a subregion of the image at time t-1 (140×140 pixels) as search image for the template which was a small crop of the image at time t (70 x 70 pixels concentric around point). Within the search image, smaller images of the size of the template were created and compared against the template using image correlation. The position with the highest similarity was the new place for the boundary point. To increase robustness of the method, the shift vectors (new position at time (t-1) – position at time t) for each point were determined and the median over the seven neighboring boundary points calculated and assigned as effective shift. Furthermore, if two boundary points were assigned to the same or neighboring pixel, one of them was removed from the computation. To ensure the quality of the annotation, we visually inspected and curated the annotation.

### Analysis of volume growth

The resulting volume measurements were analyzed using the software package R (R Core Team, 2018) using the dyplr library package (Wickham et al., 2018) for data handling. Parameters for different growth models were optimized using non-linear regression of the sum of squared differences between the actual values and the predicted value of the growth model given the parameters. As optimization algorithm, we applied a non-linear least squares approach (nls function in R). In case nls failed, we applied the Levenberg-Marquardt algorithm (nlsLM function in R, (Elzhov et al., 2016)). Plots were generated using the ggplot library (Wickham, 2016) and the gridExtra package (Auguie, 2017).

## Supporting information

## Acknowledgment

We thank all members of the Huisken lab and the Ph.D. advisory committee members Ingo Röder and Ivo Sbalzarini for constructive feedback and comments, the computer department at MPI-CBG, especially Edmund Malik and Oscar Gonzales, for providing the computer infrastructure required for this project, the fish facility at MPI-CBG (Jens Hueckmann, Joachim Hellmig, Jürgen Müller and Evelyn Lehmann) for maintaining the fish lines and the mechanical workshop at MPI-CBG (Hartmut Wolf and Falk Elsner) for producing the custom-made hardware.

## Author contributions

Stephan Daetwyler and Jan Huisken designed the project outline, carried out the experiments and analysis, interpreted results and wrote the manuscript. Ulrik Günther contributed to the segmentation and established the tools for the visualization. Carl Modes provided financial support and contributed to study design, result interpretation, and the manuscript. Kyle Harrington established the 3D segmentation pipeline and visualization, and contributed to study design, result interpretation, and the manuscript.

## Supporting Information

- S1 Detailed embedding protocol
- S2 Growth characteristics of embedded zebrafish
- S3 Sample orientation for long-term imaging of the vasculature
- S4 Automated data copying
- S5 Overview processing pipeline
- S6 Adaptations of Fiji plugins to SLURM cluster
- S7 Data stitching
- S8 Comparison of different growth models
- S9 Development of the volume of the caudalveinplexus
- S10 Phenotypic variation in zebrafish vasculature
- S11 Supplementary movies

