## Supplementary material for "Multi-sample SPIM image acquisition, processing and analysis of vascular growth in zebrafish"

### S1 Detailed embedding protocol

This supplementary chapter describes how to extend the embedding protocol established for mounting single embryos (Kaufmann et al., 2012; Weber et al., 2014) to multiple samples. Furthermore, potential difficulties and pitfalls of the embedding protocol are described and suggestion on how to solve them are mentioned.

#### 1. Preparation for multi-sample embedding

The required material for embedding was prepared (Fig. S1) and kept at room temperature for at least 5 min. In-between experiments, the embedding material (except for the FEP tubes) was stored at 4 °C.

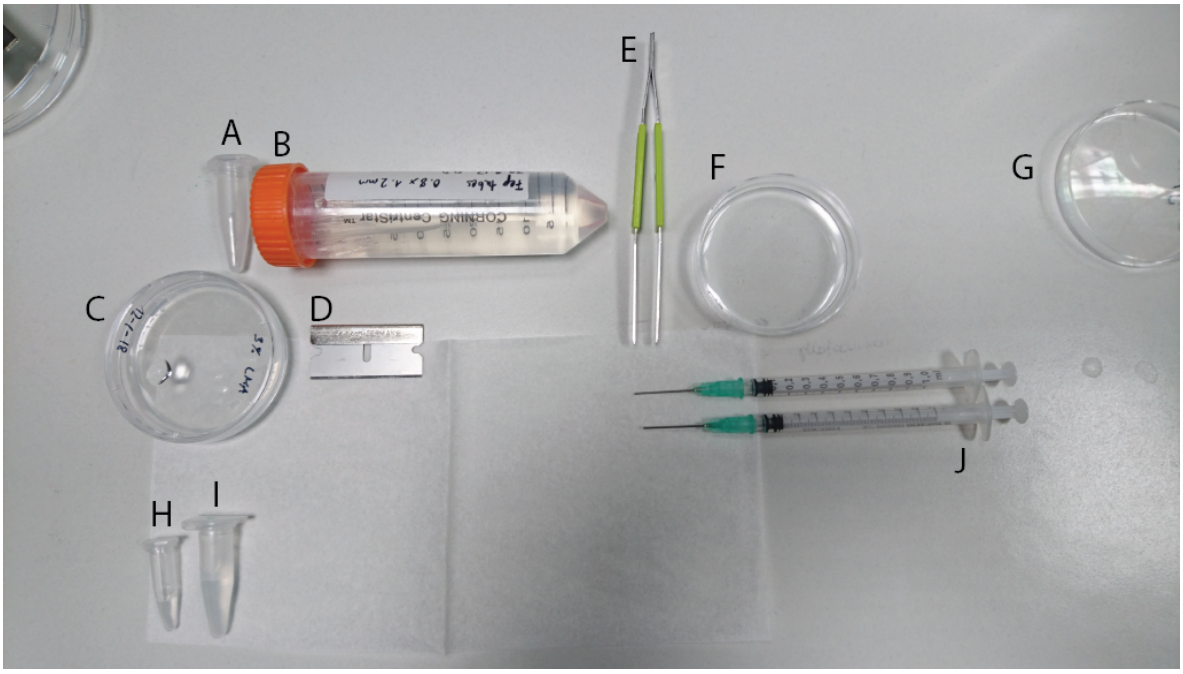


**Fig. S1: Material required for multi-sample embedding**
(A) Reaction tube containing reusable connectors (B) centrifuge tube containing cleaned FEP tubes stored in distilled water (C) dish containing a 2mm layer of 3% low melting agarose (D) razor blade (E) forceps (F) dish with prepared samples (G) dish with E3 to rinse coated FEP tubes (H) tube containing 3% methylcellulose (I) tube containing 0.1% agarose

##### Zebrafish embryos

Zebrafish (*Danio rerio*) adults and embryos were kept at 28.5 °C and handled according to established protocols (Nüsslein-Volhard and Dahm, 2002; Westerfield, 2000). The fish lines of interest were crossed into a casper background (White et al., 2008) to improve image quality by the lack of pigmentation. Male and female zebrafish were set up with a divider which was removed in the morning to time the mating of the fish. After mating, embryos were collected in a dish filled with E3 media.
The collected embryos were injected at the one or two cell stage with 2 nl of injection mix containing alpha-bungarotoxin RNA (Swinburne et al., 2015). Alpha-bungarotoxin immobilized the embryos during imaging. The injection mix was prepared of 7.7 µl distilled water, 2 µl phenolred and 0.3 µl of stock solution containing alpha-bungarotoxin RNA at a concentration of 500 ng/µl.

At around 15 hpf, the embryos expressing the fluorescent vascular marker of interest were prepared for imaging by dechorionation with sharp forceps under a stereomicroscope.

**Fluorinated Ethylene Propylene (FEP) tubes**

Fluorinated Ethylene Propylene (FEP) tubes with an inner diameter of 0.8 mm, and an outer diameter of 1.2 mm were cleaned according to established protocols (Kaufmann et al., 2012; Weber et al., 2014): First, the FEP tube were flushed with 1M NaOH and then transferred to a 50ml centrifuge tube filled with 0.5 M NaOH and ultrasonicated for 10 min. After that, the tubes were flushed with distilled water and then with 70% ethanol. After flushing the tubes, they were ultrasonicated in 70% ethanol and cut to a length of about 4 cm.
If desired, the tubes were straightened before cutting. For this, FEP tubes were inserted into steel tubes of 50 cm length. The outer diameter of the FEP tubes matched the inner diameter of the steel tube. The tubes were heated to 180 °C for 2 h in an autoclave and afterwards cooled at room temperature for at least 5 h.

**Connectors**

FEP tubes with an inner diameter of 1.1 mm and an outer diameter of 1.5 mm were used as connecting FEP tube (connector). The slightly smaller inner diameter of the connector compared to the outer diameter of the FEP tube containing the sample (1.2 mm) ensured a tight fit making use of the elasticity of FEP. The connector was prepared by mounting the FEP tube on a blunt ended needle (1.2 mm x 40 mm). On this, a hole of about 2 mm was cut in the middle of the connector and the connector cut to its final length of about 6 mm. The connectors were reused in several experiments.

**3% methylcellulose for coating**

3% methylcellulose was prepared for coating the FEP tube to prevent attachment of the growing fish to the FEP wall. E3 was heated to approximately 60-70 °C and methylcellulose powder was added to a final concentration of 3%. To ensure a homogenous solution, it was stirred at 4 °C overnight.

**0.1% agarose in E3 and 3% agarose dish for plugging**

The appropriate amount of agarose was dissolved in E3 and heated up in a microwave until the agarose solution appeared homogeneous. 1.2 ml aliquots of 0.1% agarose were prepared in 1.5 ml reaction tubes and stored at 4 °C. Similarly, the 3% low melting agarose was poured into small petri-dishes (60 mm x 15 mm) to a thickness of the low melting agarose layer of about 2 mm. After solidifying, E3 medium was added on top of the low melting agarose layer to prevent it from drying. Also the petri-dish with the low melting agarose was stored at 4 °C.

#### 2. Embedding of single samples

To embed multiple embryos, the protocol for single sample embedding (Kaufmann et al., 2012; Weber et al., 2014) was followed. The use of gloves during the embedding procedure prevented contamination and impairment of imaging by finger prints on the tubes. Furthermore, bending the FEP tubes at this stage was avoided as the FEP tube might relax to its previous state during time-lapse imaging and consequently move the sample out of the field of view.

**Coating of the FEP tube**

First, the FEP tube was coated with 3% methylcellulose. For this, the FEP tube (0.8 mm I.D./1.2 mm O.D.) was attached to a syringe and the tube filled with methylcellulose. The methylcellulose was then removed from the tube so that only a thin coating remained on the tube wall.

**Mounting of zebrafish embryos**

First, the coated FEP tube was prepared for sample mounting by filling it with 0.1% agarose. Using a glass Pasteur pipette, a selected zebrafish embryo was transferred from the petri dish to the reaction tube containing 0.1% agarose at room temperature. Ideally, as little additional E3 was transferred with the zebrafish to the reaction tube.
Using the syringe, the embryo was then pulled very gently into the tube towards the tip of the needle with its head first. Repeated pulling and pushing of the embryo was avoided. With a sharp razor blade the tube was cut 1 to 2 mm above the head of the zebrafish embryo.
The FEP tube with the sample was then plugged with 3% low melting agarose. Before plugging the FEP tube, it was tipped slightly several times on the 3% low melting agarose dish so that only a minimal distance between head and plug remained after embedding. It was however avoided that the embryo touched the 3% melting agarose layer. After that, the tube was placed straight on the 3% low melting agarose layer and turned slowly (like a screw) into the 3% plugging layer.
After embedding, the embryo was checked for integrity under the stereoscope.

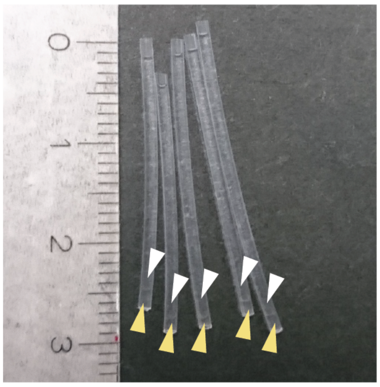

**Fig. S2: Mounted single sample embryos**
After mounting, zebrafish embryos (white arrowhead) were resting on a 3% low melting agarose plug (yellow arrowhead)

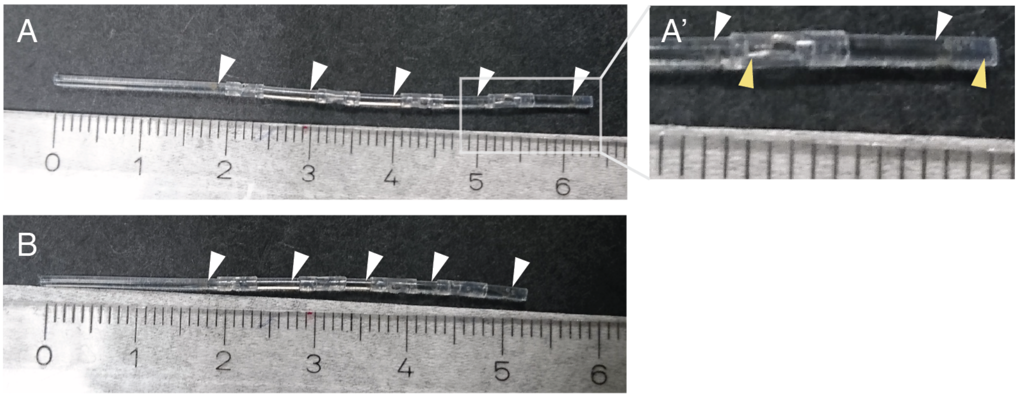


**Fig. S3: Mounting of several embryos**
Several embryos (white arrowheads) were mounted together for one experiment. (A) The connectors were stacked so that the whole embryo was accessible (A’) Zoom in to A containing two fish (white arrowheads) and two plugs (yellow arrowheads) (B) only the head of the embedded fish was accessible resulting in a shorter overall tube.

#### 3. Preparation of the multi-sample tube

The FEP tubes containing one mounted zebrafish embryos (Fig. S2) were cut to a length of about 9 mm providing space for the plug (about 2-3 mm), the zebrafish embryo (at the end of

the time-lapse around 4 mm), and space for the connector (about 2-3 mm). The individual tubes were stacked together to form one long tube (Fig. S3). Depending on the experimental requirements, the overlap of the connector with the FEP tubes containing the sample could be varied. If only parts of the embryo such as the head were imaged, the tubes could be stacked with a higher overlap resulting in an overall shorter tube or the possibility to image more samples (Fig. S3, B).
The hole in the tube not only increased oxygen availability in the sample tube but also made the stacking easier as no air pressure built up when stacking the tubes together.

#### 4. Potential embedding difficulties

Mounting of zebrafish embryos can be challenging. Below, some pitfalls and suggestions how to solve them are listed (Table S1)

**Table S1: Suggestions for potential embedding problems**

| Problem | Solution |
|---|---|
| Dying embryos | It is important that the embedding material is at room temperature before embedding. Zebrafish growth and survival is affected by too cold and too warm temperatures.<br>After embedding, the integrity of the embryo should be checked under the stereomicroscope. Ideally, the embryo lies just above the plug but does not touch it yet.<br>If dying samples reoccur, also changing the companies providing the materials should be considered. It has been reported orally that FEP tubes from some companies might be more likely to decrease sample health. The same applies for agarose. Also check for pH and possible contamination of E3. We only applied filtered E3. |
| Formation of air bubbles | To avoid air bubbles, empty the syringe after sucking up the methylcellulose by disconnecting it from the cannula. Put the syringe and cannula back together, wipe the methylcellulose from the exterior of the FEP tube and rinse it several times with E3. Before mounting the embryo in the tube, also suck up some 0.1% agarose.<br>Moreover, fill the dish containing the 3% low melting agarose to produce the plug with a bit of E3. With this, formation of air bubbles during plugging is reduced.<br>The 0.1% agarose might contain air bubbles. To remove them, vortex the reaction tube shortly before putting the fish into it. |
| Zebrafish is far away from the plug after embedding | Before plugging, tap the zebrafish several times on the 3% low melting agarose without plugging. This produces a suction force if the FEP tube is long enough. A length of 3.5 – 5 cm showed to be ideal for this. |
| Zebrafish moves during imaging | If you apply Tricaine as described in the original sample mounting paper (Kaufmann et al., 2012), zebrafish move at around 1 dpf. Therefore, we used alpha-bungarotoxin RNA in our long-term imaging studies. Also make sure that the embryo is close to the plug before starting an imaging experiment and that the FEP tubes have not been bent before the imaging was started. |

#### 5. Material list

**Table S2: List of Materials required for embedding**

| **Name** | **Company / Catalog Number** | **Comments** |
|---|---|---|
| Agarose, low melting temperature | Sigma / A9414 | Supplied as powder |
| Bungarotoxin RNA | | Stocks of 500 ng/ul were generated as described in (Swinburne et al., 2015) |
| Greiner Petri Dish | Sigma | Plastic, 94 mm x 16 mm and 60x15 mm |
| E3 | | Prepared as described in (Nüsslein-Volhard and Dahm, 2002), p. 22 |
| Fluorinated Ethylene Propylene (FEP) tubes | Fluidflon FEP-Schlauch, Liquid Scan, pro liquid GmbH (http://www.proliquid.de) Art: 2001048_E | For the tube containing the growing zebrafish, an inner diameter (I.D.) of 0.8 mm and an outer diameter (O.D.) of 1.2 mm was chosen.<br>The connectors were made of FEP tube with an I.D. of 1.1 mm and an O.D. of 1.5 mm. |
| Forceps | DUMONT / Dumostar Nr. 55 – Article 11295-51 | Used for decorionation of zebrafish embryos |
| Methylcellulose | Sigma / M0387 | Supplied as powder |
| Omnifix-F Solo 1 ml Syringe | B. Braun Melsungen AG / 9161406V | |
| Phenol red solution | Sigma / P0290 | For injection mix |
| Sterican Single-Use Cannula, blunt | B. Braun Melsungen AG Sterican 21G x 7/8’’<br>BD Blunt Fill Needle, 18G 1 1/2 | For embedding, 0.8 mm x 22 mm cannula were used. To prepare the connectors, blunt ended needles (1.2 mm x 40 mm) were used. |
| Tricaine / MS-222 | Sigma / E10521 | Alternative mean of zebrafish immobilization instead of using bungarotoxin |
| 70% EtOH | | To clean the tube |
| 1 M NaOH | Merck, 1091371000 | To clean the tube |
| 50 ml Polypropylene Centrifuge Tube | Corning / 430829 | 15 ml and 50 ml tubes were used as storage containers |
| 1.5 ml reaction tube | Eppendorf / 3810X | |

### S2 Growth characteristics of embedded zebrafish

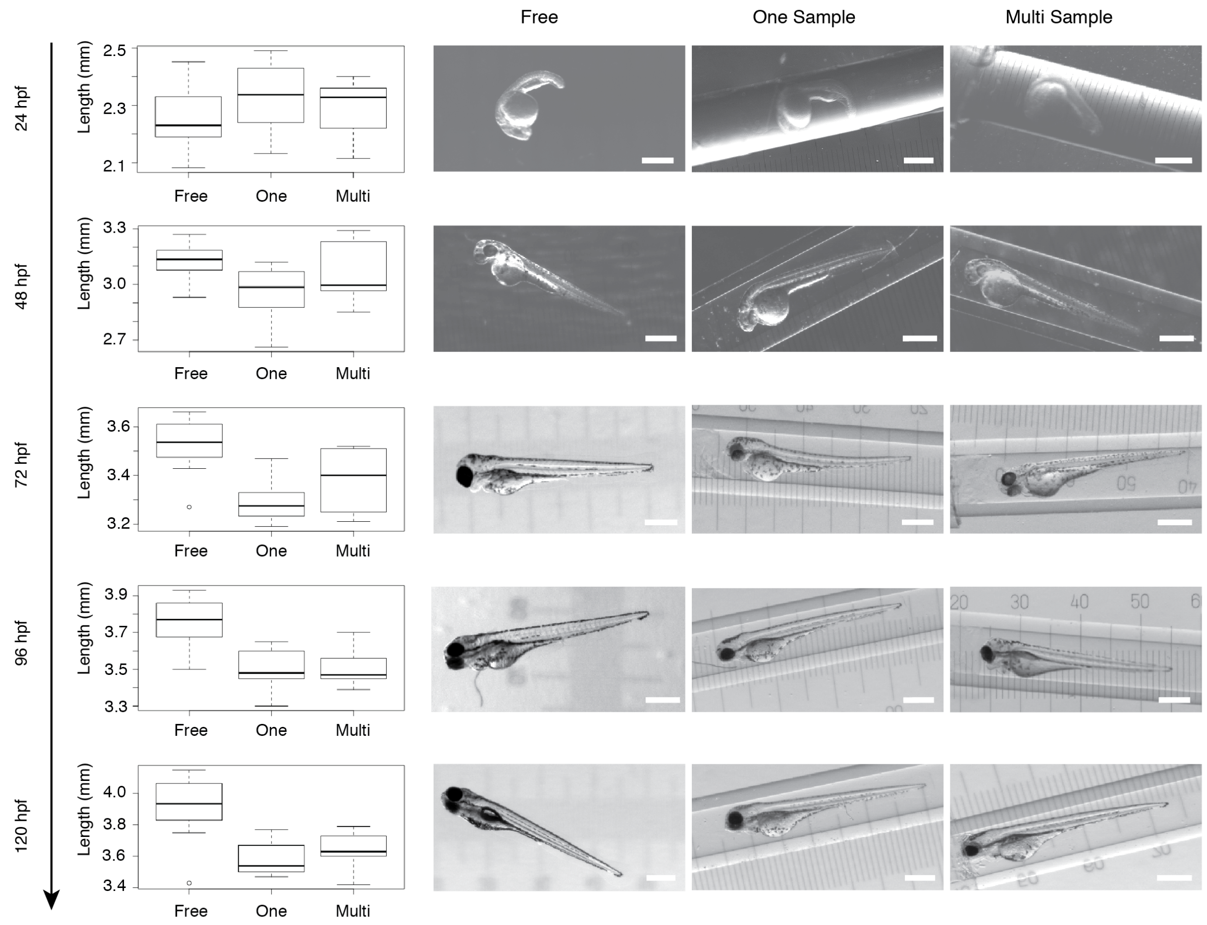


**Fig. S4: Multi-sample embedding**

Distribution of body length measurements of freely swimming fish (n=8), single embedded samples (n=10), and samples embedded in the multi-sample tube (n=9) with representative images for each embedding. The fish were embedded at 24 hpf. No morphological defects were observed with all three embedding protocols. Scale bar 500 μm.

### S3 Sample orientation for long-term imaging of the vasculature

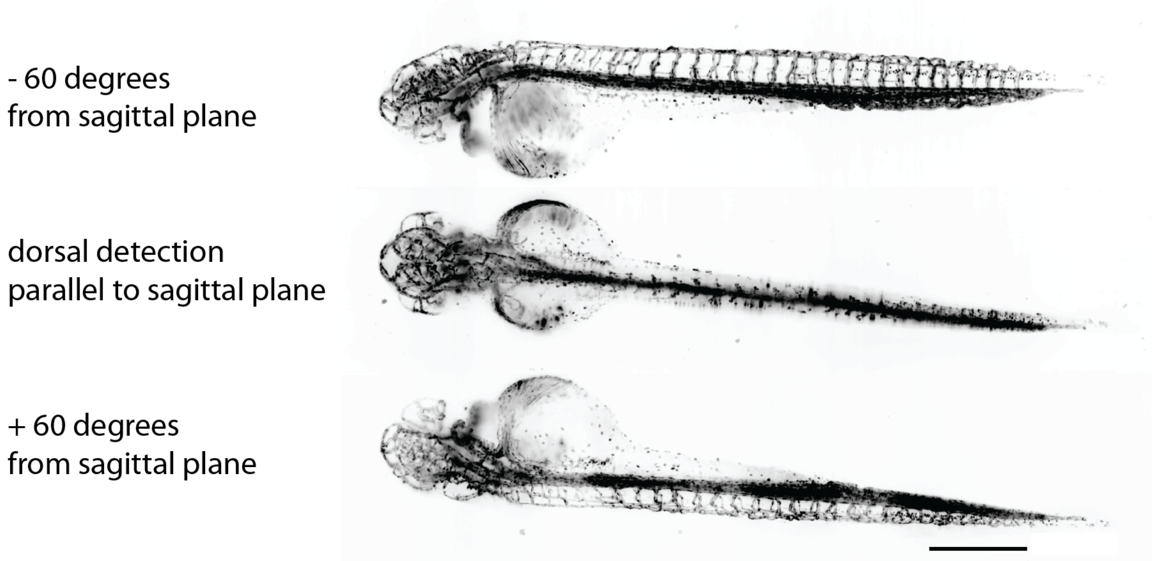


**Fig. S5: Imaging directions of zebrafish.**
Zebrafish embryos were imaged from three direction to have full access to the whole vasculature. Scale bar: 0.5 mm.

### S4 Automated data copying

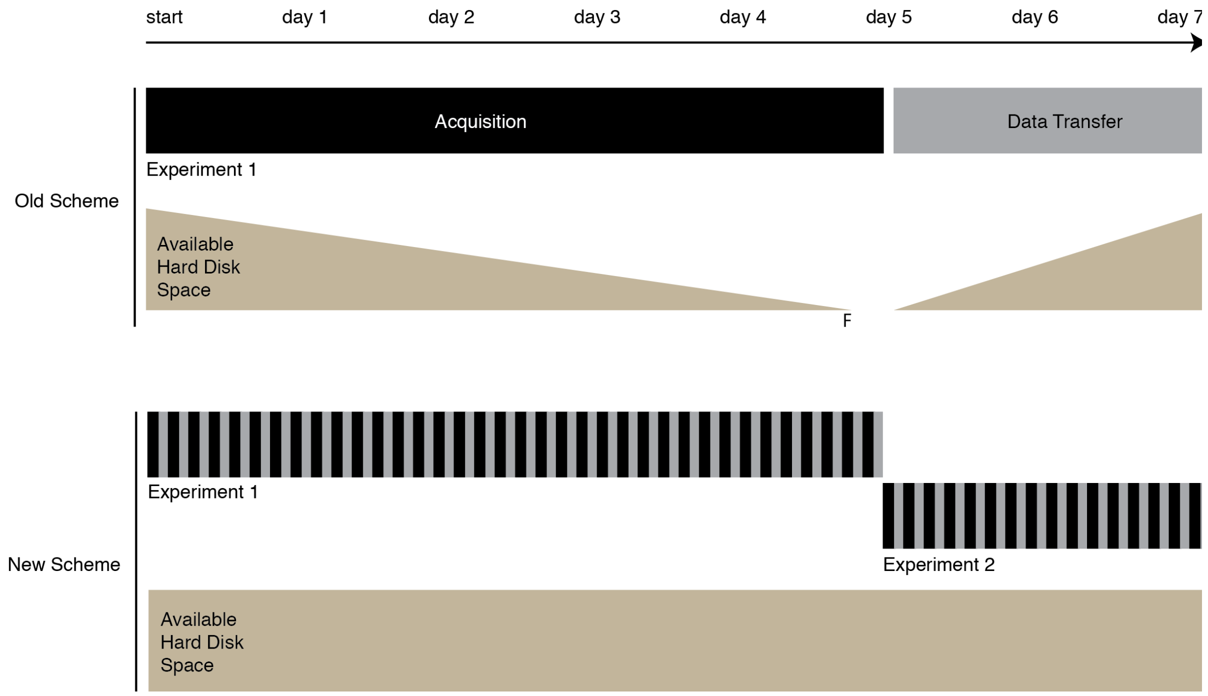


**Fig. S6: Data transfer during acquisitions.**
Comparison of old and new scheme. In the old scheme, data acquisition (black) and transfer (gray) were sequential. With this acquisition scheme, there was a risk of filling the hard disks (F) before the end of the planned acquisition time. In the new scheme, after every acquisition (black), data is transferred (gray). Ideally, all data acquired was transferred, and therefore the available hard disk space did not decrease. Moreover, already a new experiment could be started at the end of first experiment leading to higher throughput.

#### S5 Overview processing pipeline

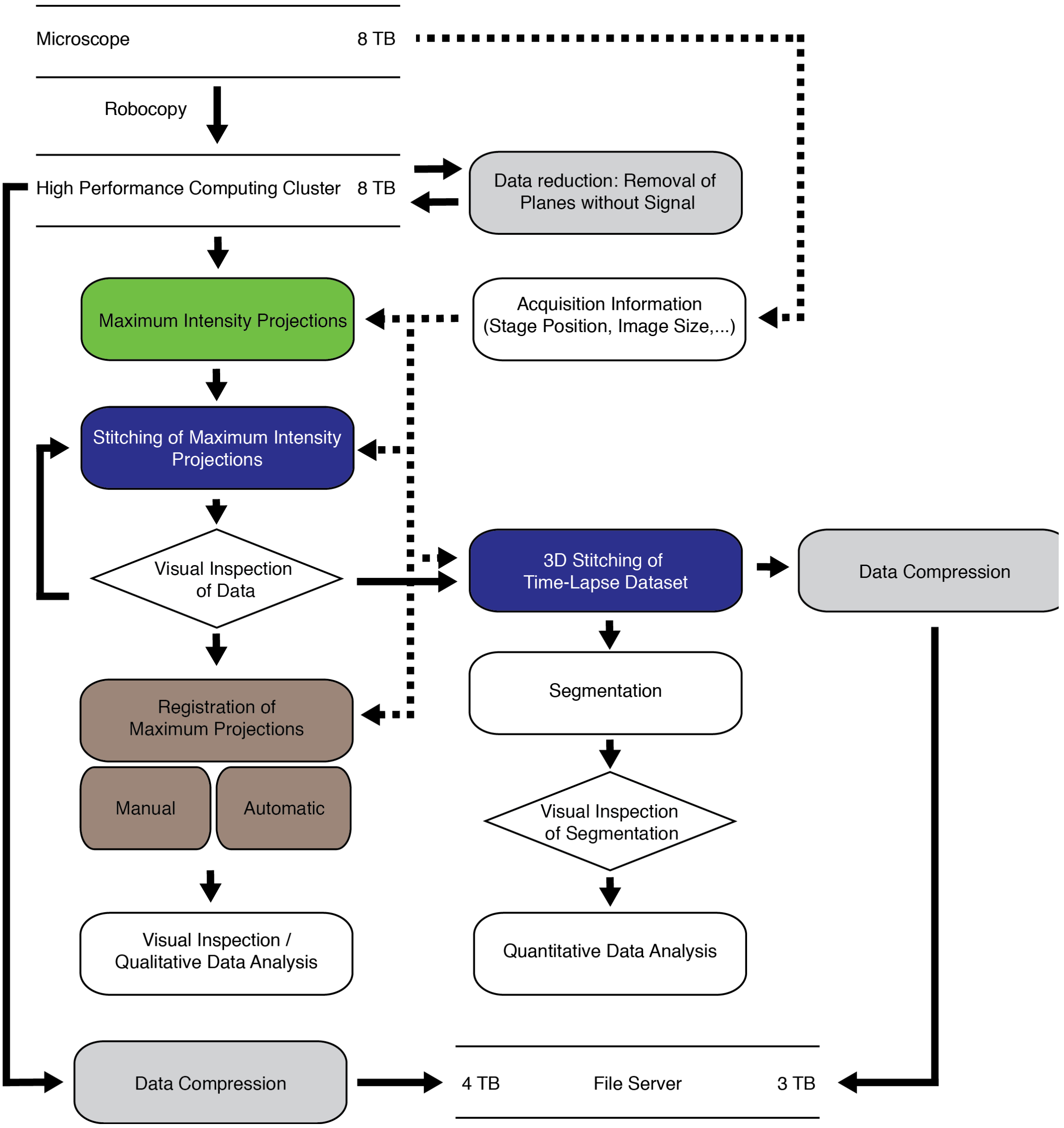


**Fig. S7: Overview about data processing pipeline**
The dedicated data processing pipeline was composed of modular parts for data compression (grey), projections (green), stitching of the data (blue), registration (brown), analysis (white). Full arrows indicated data flow, dashed arrow indicated the input of parameters from the microscope such as stage positions to the processing steps. Data quality was evaluated visually (◊) to ensure proper processing of the data.

### S6 Adaptation of Fiji plugins to SLURM cluster

1) To run the Fiji plugin on the cluster, add code so that macro input can be parsed on the cluster without running the GUI of FIJI.

```
import ij.IJ;
import ij.ImageJ;
import ij.Macro;
import ij.gui.GenericDialog;
import ij.plugin.PlugIn;

public class SPIM_yourplugin_ implements PlugIn {

	@Override
	public void run(String args) {

		//generate an object of GenericDialog class to which options can be
		//added and which can be evaluated
		final GenericDialog gd = new GenericDialog( "Indicate folder" );

		//generate options for the Plugin to work on SPIM data from the
		//multi-sample SPIM
		gd.addStringField( "Foldername ", "D:\\processfolder", 50 );
		gd.showDialog();

		if ( gd.wasCanceled() )
			return;

		// read in input from plugin interface
		String directory_name = gd.getNextString();

		// read in input if run as a plugin/macro on the cluster
		if(IJ.isMacro()){
			String options = Macro.getOptions();
			String[] listOptions = options.split(" ");
			String[] parsedOptions = new String[listOptions.length];
			IJ.log(options);
			for(int iter=0; iter<listOptions.length; iter++){
				String iterString = listOptions[iter];
				String[] tmplist = iterString.split("=");
				IJ.log(iterString);
				parsedOptions[iter] = tmplist[1];

			}
			directory_name = parsedOptions[0];
		}

		IJ.log(directory_name);
	}

public static void main(String[] args2) {

		new ImageJ();
		IJ.runPlugIn("SPIM_yourplugin_ ","");
}
}
```

2) Install Fiji on the cluster, e.g. to the folder projects/yourproject/fiji

3) To launch the plugin on a cluster, here running with SLURM, it is convenient to prepare two files: (i) interface with SLURM (ii) interface with Fiji

(i) File to interface with slurm, e.g save as "start_yourplugin.sh"

```
#!/bin/bash
#SBATCH --nodes=1
#SBATCH --tasks-per-node=24
#SBATCH --cpus-per-task=1
#SBATCH --time=16:00:00
#SBATCH --mem 62000

/projects/yourproject/fiji/Fiji.app/ImageJ-linux64 --headless -macro
./run_yourplugin.ijm
```

Note: Here you specify parameters for the cluster, such as the number of nodes, tasks-per-node, CPUs per task, the time you estimate your plugin to run and the memory requirements for the tasks.

(ii) File to interface with Fiji, e.g. save as "run_yourplugin.ijm"

```
//set parameters for renaming and copy files
Foldername = "/projects/yourfolder";

//run it
run("SPIM_yourplugin_", "foldername =" +Foldername);
```

Note: The name of the plugin here `"SPIM_yourplugin_"` is the name specified in the scr/main/resources, plugins.config file.

(iv) now you can start your Fiji plugin from the command line:

```
sbatch start_yourplugin.sh
```

Note: If the .sh file is not found, be sure to be in the right folder where you also saved your .sh file

### S7 Data stitching

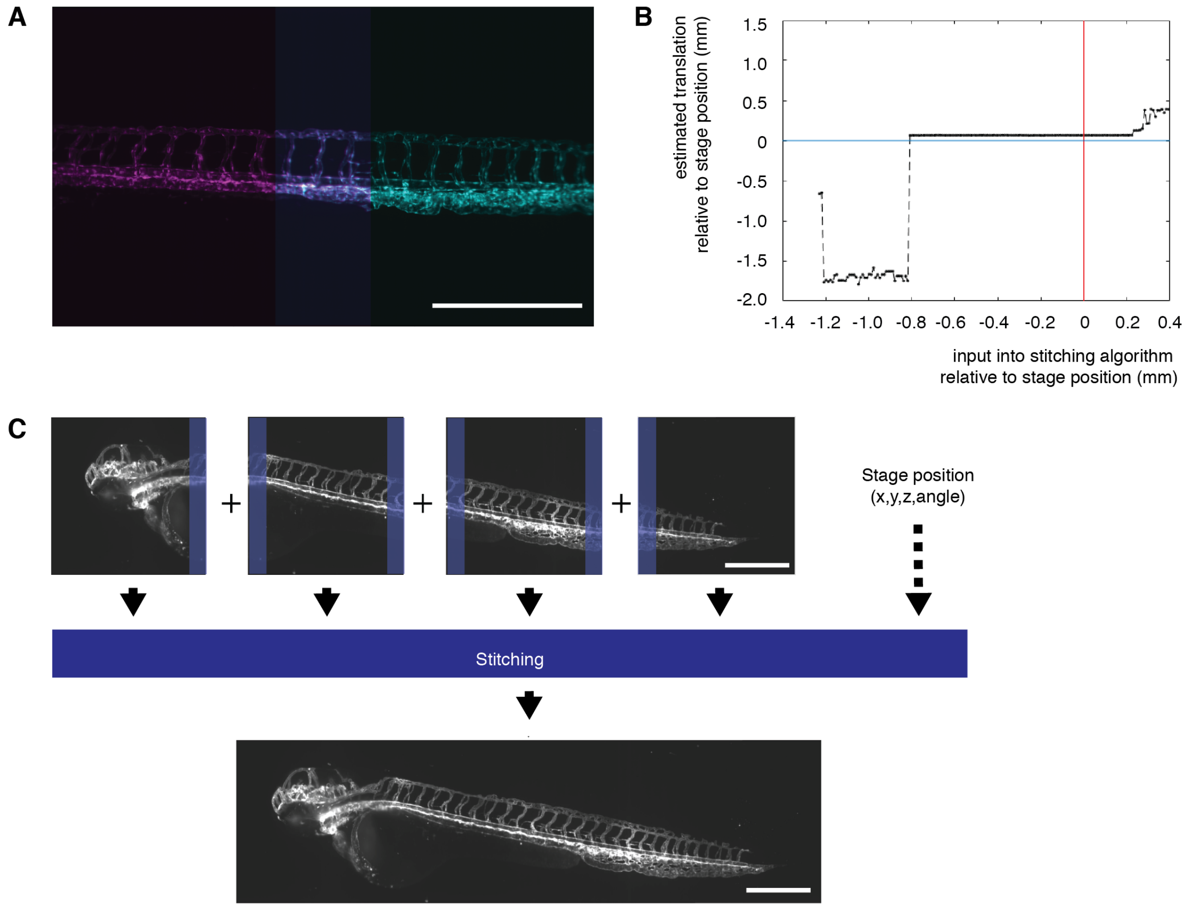


**Fig. S8: Combining multiple acquisition volumes into one image: data stitching.**
(A) Stitching of two images (magenta and cyan) zebrafish expressing the vascular marker Tg(kdrl:EGFP) (Jin et al., 2005). The overlap between the two images is highlighted in violet. (B) To automatically calculate the stitching of the two images in (A), an initial offset was set to initialize the calculation. Varying this offset revealed that it was robust over a large range of initialization values, nevertheless, stitching became unreliable if the value were too far off from the assumed offset given by the translational stages (red line). Moreover, the stitching result indicated that the calculated translation (black line) was close to the physical coordinate given by the stage position (blue line) but not exact (0.06 mm / 53 pixels away). Therefore, using the stage position was a reasonable estimate for initialization. (C) In my implementation of stitching, stage positions determined which tiles to combine and then based on the initialization, the stitching generated a final output image. Here, a zebrafish expressing the vascular marker Tg(kdrl:Hsa.HRASmCherry) (Chi et al., 2008) was displayed . Scale bar 0.5 mm.

### S8 Comparison of different growth models

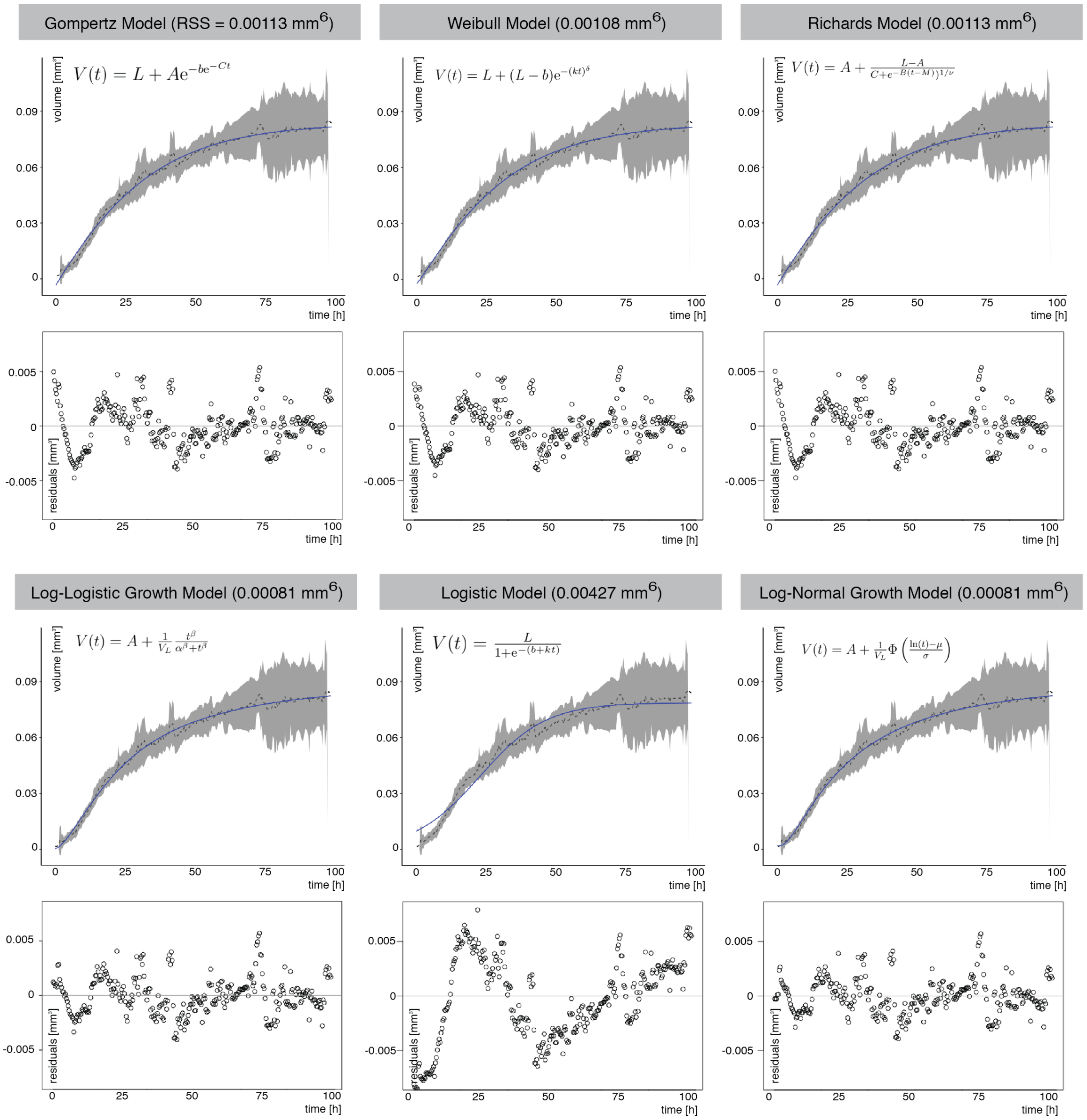


**Fig. S9: Comparison of different growth models**

Different growth models were compared and their residual sum of errors (RSS) calculated. A low RSS indicated a good fit. For each growth model, the predicted values from the model (blue line) were overlaid with the experimental data (dashed line – mean, gray ribbon 95% confidence interval). Moreover, for each growth model, the residuals were plotted over time. Small residual values combined with a random distribution of them around the 0 value (gray line) indicated good description of the data by the growth model. The log-normal and log-logistic growth models represented the data the best of all the tested models.

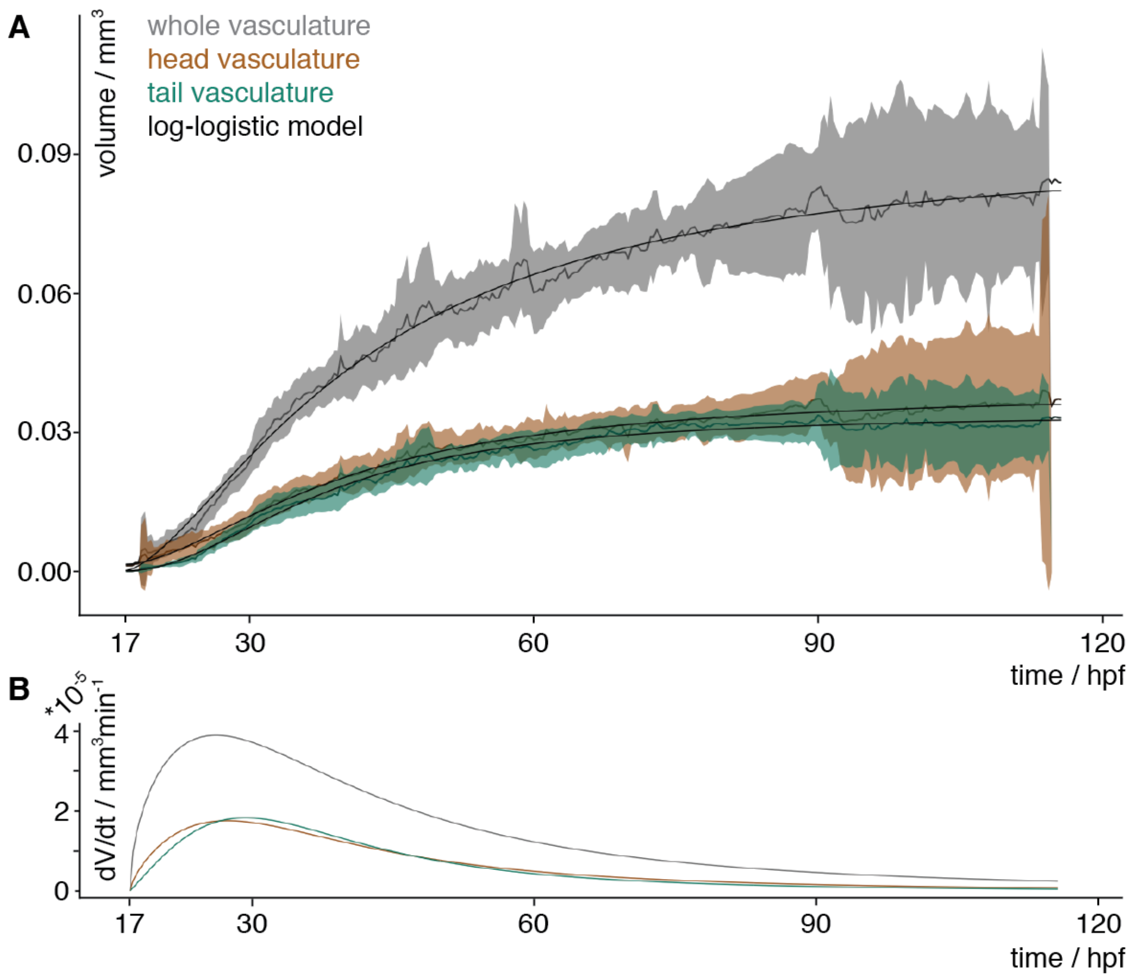


**Fig. S10: Vascular volume growth characteristics of zebrafish using the log-logistic data**
**(A)** Experimental measurements of the volume over time of the whole vasculature (gray), the head (brown) and tail (turquoise) vasculature. The mean of the measurements is depicted with a solid line and the 95% confidence interval (t-statistics, n=7) as a ribbon in the corresponding color. The black line depicts the approximation of the volume by the log-logistic growth model. (**D**) The volume growth rate of the whole vasculature (gray), head (brown) and tail (turquoise) was calculated by inserting the parameters obtained from the approximation into the change of volume equation from (B).

#### S9 Development of the volume of the caudalveinplexus

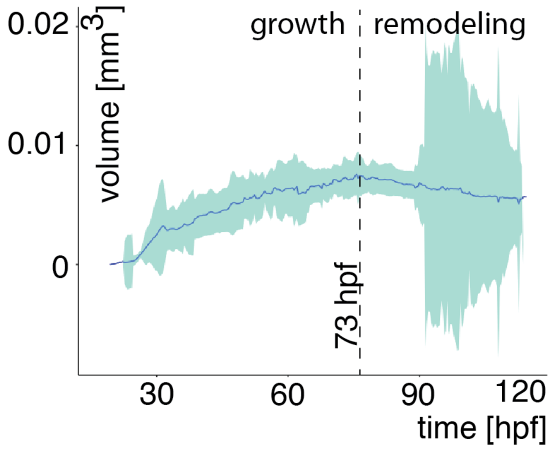


**Fig. S11: Log-logistic growth characteristics**
The volume of the caudalveinplexus in the tail grew until around 73 hpf after which its size started to decrease. The mean of the measurements is depicted with a blue line and the 95% confidence interval (t-statistics, n=7) as a ribbon in the light turquoise.

#### S10 Phenotypic variation in zebrafish vasculature

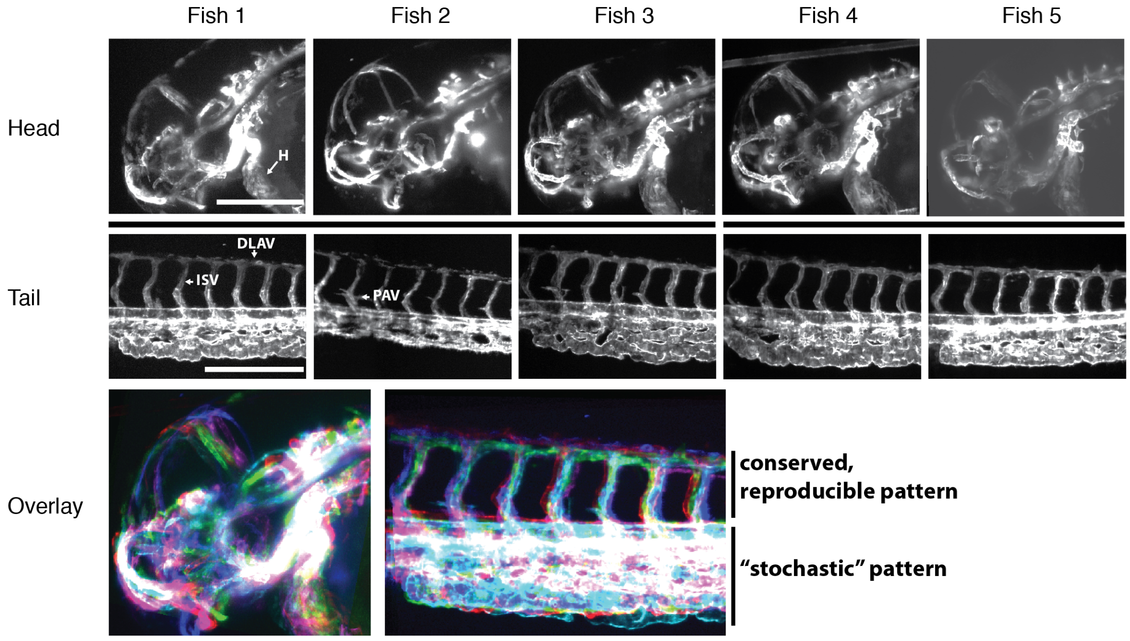


**Fig. S11: Phenotypic variation of zebrafish vasculature**
Five zebrafish expressing the vasculature marker Tg(kdrl:lifeactGFP) (Vanhollebeke et al., 2015) at 35 hpf were imaged using SPIM. Maximum projections of the head and tail of five different individual zebrafish embryos (H, heart; DLAV, dorsal longitudinal anastomotic vessel; ISV, intersegmental vessels; PAV, parachordal vessel). Scale bar: 0.25 mm. Overlay of the head and the tail parts revealed different degrees of variation in the vasculature. The bright white signal indicated maximum overlap.

### S11 Supplementary movies

##### Movie 1

Zebrafish embryos expressing the fluorescent vascular marker *Tg(kdrl:EGFP)* in cyan and the red blood cell marker, *Tg(GATA1a:dsRed)* in magenta imaged from three different angles over several days. Scale bar: 0.5 mm.

##### Movie 2

Three different zebrafish embryos expressing the fluorescent vascular marker *Tg(kdrl:EGFP)* in cyan and the red blood cell marker, *Tg(GATA1a:dsRed)* in magenta imaged in the same experiment simultaneously over several days. Scale bar: 0.5 mm.

##### Movie 3

3D-rendered segmentation of the zebrafish vasculature with annotations of the head vasculature in orange, the tail in turquoise with its caudalveinplexus highlighted with light-turquoise and unannotated vasculature in grey.

**Chi, N. C., Shaw, R. M., De Val, S., Kang, G., Jan, L. Y., Black, B. L. and Stainier, D. Y. R.** (2008). Foxn4 directly regulates tbx2b expression and atrioventricular canal formation. *Genes Dev* **22**, 734–739.

**Jin, S.-W., Beis, D., Mitchell, T., Chen, J.-N. and Stainier, D. Y. R.** (2005). Cellular and molecular analyses of vascular tube and lumen formation in zebrafish. *Development* **132**, 5199–5209.

**Kaufmann, A., Mickoleit, M., Weber, M. and Huisken, J.** (2012). Multilayer mounting enables long-term imaging of zebrafish development in a light sheet microscope. *Development* **139**, 3242–3247.

**Nüsslein-Volhard, C. and Dahm, R.** (2002). *Zebrafish: A Practical Approach*. Oxford University Press.

**Swinburne, I. A., Mosaliganti, K. R., Green, A. A. and Megason, S. G.** (2015). Improved Long-Term Imaging of Embryos with Genetically Encoded α-Bungarotoxin. *PLoS ONE* **10**, e0134005.

**Vanhollebeke, B., Stone, O. A., Bostaille, N., Cho, C., Zhou, Y., Maquet, E., Gauquier, A., Cabochette, P., Fukuhara, S., Mochizuki, N., et al.** (2015). Tip cell-specific requirement for an atypical Gpr124- and Reck-dependent Wnt/β-catenin pathway during brain angiogenesis. *Elife* **4**, 2807.

**Weber, M., Mickoleit, M. and Huisken, J.** (2014). Multilayer Mounting for Long-term Light Sheet Microscopy of Zebrafish. *J. Vis. Exp.* e51119–e51119.

**Westerfield, M.** (2000). *The zebrafish book. A guide for the laboratory use of zebrafish Danio rerio*. 4 ed. Eugene: Univ. of Oregon Press.

**White, R. M., Sessa, A., Burke, C., Bowman, T., LeBlanc, J., Ceol, C., Bourque, C., Dovey, M., Goessling, W., Burns, C. E., et al.** (2008). Transparent adult zebrafish as a tool for in vivo transplantation analysis. *Cell Stem Cell* **2**, 183–189.
